# Synthetic Biology Tool Development Advances Predictable Gene Expression in the Metabolically Versatile Soil Bacterium *Rhodopseudomonas palustris*

**DOI:** 10.1101/2021.11.01.466785

**Authors:** Cheryl Immethun, Mark Kathol, Taity Changa, Rajib Saha

## Abstract

Harnessing the unique biochemical capabilities of non-model microorganisms would expand the array of biomanufacturing substrates, process conditions, and products. There are non-model microorganisms that fix nitrogen and carbon dioxide, derive energy from light, catabolize methane and lignin-derived aromatics, are tolerant to physiochemical stresses and harsh environmental conditions, store lipids in large quantities, and produce hydrogen. Model microorganisms often only break down simple sugars and require low stress conditions, but they have been engineered for the sustainable manufacture of numerous products, such as fragrances, pharmaceuticals, cosmetics, surfactants, and specialty chemicals, often by using tools from synthetic biology. Transferring complex pathways with all of the needed cofactors, energy sources, and cellular conditions from a non-model microorganism to a common chassis has proven to be exceedingly difficult. Utilization of unique biochemical capabilities could also be achieved by engineering the host; although, synthetic biology tools developed for model microbes often do not perform as designed in other microorganisms. The metabolically versatile *Rhodopseudomonas palustris* CGA009, a purple non-sulfur bacterium, catabolizes aromatic compounds derived from lignin in both aerobic and anaerobic conditions and can use light, inorganic, and organic compounds for its source of energy. *R. palustris* utilizes three nitrogenase isozymes to fulfill its nitrogen requirements while also generating hydrogen. Furthermore, the bacterium produces two forms of RuBisCo in response to carbon dioxide/bicarbonate availability. While this potential chassis harbors many beneficial traits, stable heterologous gene expression has been problematic due to its intrinsic resistance to many antibiotics and the lack of synthetic biology parts investigated in this microbe. To address these problems, we have characterized gene expression and plasmid maintenance for different selection markers, started a synthetic biology toolbox specifically for the photosynthetic *R. palustris*, including origins of replication, fluorescent reporters, terminators, and 5’ untranslated regions, and employed the microbe’s endogenous plasmid for exogenous protein production. This work provides essential synthetic biology tools for engineering *R. palustris*’ many unique biochemical processes and has helped define the principles for expressing heterologous genes in this promising microbe through a methodology that could be applied to other non-model microorganisms.

## Introduction

Traditional abiotic chemical synthesis can be energy intensive, create toxic byproducts that have a deleterious effect on the environment, and require a multitude of steps to create the often complex and chiral compounds (Wang and Weller, 2006;Engels et al., 2008;Ajikumar et al., 2010;Sasidharan et al., 2011). Processes utilizing microorganisms can address many of these problems while taking advantage of unique biochemical reactions often found in non-model bacteria (Basak and Das, 2007;Yan and Fong, 2017;Liang et al., 2020). For instance, various *Geobacter* species can produce bioelectricity (Lovley et al., 2011;Rosenbaum and Henrich, 2014). *Burkholderia sacchari* (Guamán et al., 2018), *Psuedomonas spp*., and *Halomonas spp*. (Zhang et al., 2020) synthesize polyhydroxyalkanoates (PHAs), sustainable alternatives to petroleum-derived plastics. Many purple non-sulfur bacteria can also produce PHAs as well as hydrogen, and catabolize a wide array of substrates that are commonly considered waste products (Basak and Das, 2007;Özgür et al., 2010;Brown et al., 2020;Alsiyabi et al., 2021).

Model organisms like *Escherichia coli* are most commonly used in biochemical processes due to their rapid growth rate and comprehensive characterization. *E. coli* has been engineered to produce a number of value-added chemicals, including ethanol (Shams Yazdani and Gonzalez, 2008), 1-butanol (Atsumi et al., 2008) methyl-ketone (Wang et al., 2018), a styrene monomer (Liang et al., 2020), and various pharmaceuticals from organic feedstocks (Chang and Keasling, 2006;Kizer et al., 2008). However, many attempts to express heterologous pathways in model bacteria have proven to be fraught with difficulties. In addition to the substantial engineering work required to introduce all of the edits into the genome, synthesis of all of the new proteins causes a severe increase in the host’s metabolic load and subsequent growth deficits (Kizer et al., 2008;Müller et al., 2015). Furthermore, the new pathway can produce metabolites that are toxic to non-native host (Chang and Keasling, 2006;Kizer et al., 2008;Nowroozi et al., 2014;Müller et al., 2015;Ng et al., 2015). Model microorganisms can also lack the necessary metabolites to synthesize the desired products (Müller et al., 2015;Wu et al., 2018). In addition, the pathway can be imbalanced in the new host, leading to a shortage of required enzymes for crucial reactions (Nowroozi et al., 2014). This can lead to lower product yields than what was found in the native organism (Wu et al., 2018). Engineering the native host to harness its vast metabolic abilities as the natural producer of the value-added chemical(s) could address these issues.

To harness the large metabolic potential of non-model bacteria, it is vital to build tools that enable precise and predictable gene expression in these microorganisms. Synthetic biology toolkits have been developed for non-model organisms such as cyanobacteria in order to increase titers of value-added chemicals these organisms produce (Markley et al., 2015;Nozzi et al., 2017;Mukherjee et al., 2020). Cyanobacteria’s ability to conduct photosynthesis has also been utilized for the production of chemicals such as ethanol (Namakoshi et al., 2016), ethylene (Wang et al., 2018), and isoprene (Bentley et al., 2014). Toolkits have also been developed for thermophiles for the production of organic acids and biofuels like ethanol, hydrogen, and butanol (Zeldes et al., 2015). These toolkits include methods to regulate gene expression in non-model organisms such as RBS optimization (Ng et al., 2015;Nieuwkoop et al., 2019), organism specific promoter and terminator libraries (Du et al., 2012;Elmore et al., 2017), and context optimization, since the order of the genes affects transcription rates (Smanski et al., 2014). Despite this progress, the development of synthetic biology tools for non-model organisms is still limited, as the performance of genetic parts are often organism specific (Martínez-García and de Lorenzo, 2017;Yan and Fong, 2017).

*Rhodopseudomonas palustris* CGA009 (hereafter *R. palustris*) is a purple non-sulfur bacterium (PNSB) capable of all four modes of metabolism. *R. palustris*’ chemotrophic and phototrophic abilities provide the energy necessary for energy-intensive biochemical reactions, such as nitrogen and carbon fixation (Larimer et al., 2004). *R. palustris* can catabolize a variety of carbon sources, including a vast array of aromatic compounds such as lignin breakdown products (Harwood and Gibson, 1988;Ramasubramanian et al., 1996;Barbosa et al., 2001;Austin et al., 2015). It can also produce polyhydroxybuturate (Brown et al., 2020) and hydrogen (Barbosa et al., 2001;Huang et al., 2010), and can store up to 39% of its biomass as fatty acids that can be used in biofuels (Carlozzi et al., 2010). *R. palustris* has also gained interest as a tool for bioremediation as it possesses the ability to remove nutrients from wastewater to within European standards (Cerruti et al., 2020). All of *R. palustris*’ beneficial traits make the metabolically robust organism an ideal candidate as a biotechnology chassis. Extensive research has been done on the characterization and improvement of the hydrogen production (Gosse et al., 2010;McKinlay et al., 2014) and the PHB production capabilities of *R. palustris* (Ranaivoarisoa et al., 2019;Alsiyabi et al., 2021). In addition, *R. palustris* has been engineered to produce n-butanol (Doud et al., 2017;Bai et al., 2020). Despite the engineering interest in *R. palustris*, their success has been limited.

Several difficulties inhibit the use of *R. palustris* as a biocatalyst. Engineering efforts with *R. palustris* struggle with its natural resistance to antibiotics (Larimer et al., 2004), requiring high concentrations to maintain selective conditions (Welander et al., 2012;Xu et al., 2016). Transient expression of genes expressed from non-native plasmids has also been a problem (du Toit et al., 2021). In addition, genetic parts essential for engineering the bacterium have not been tested in *R. palustris*. Fluorescent reporters enable in vivo characterization of synthetic biology parts; yet, the background fluorescence can complicate their use in pigmented bacteria like *R. palustris*. The performance of basic building blocks of gene expression, such as origins of replication, 5’ untranslated regions (UTRs), and transcriptional terminators, has also not been determined for this microbe, limiting the biotechnology applications. This study tackles these problems by characterizing the bacterium’s *‘famous’* insensitivity to antibiotic selection, the stability of exogenous plasmids, and the performance of fluorescent reporters. In addition, two transcriptional terminators are tested to minimize the effect of genetic context, and design rules for 5’ UTRs are explored. To improve the maintenance of heterologous gene expression in *R. palustris*, an expression cassette was then integrated into three sites in the bacterium’s endogenous plasmid. Adding these tools for producing heterologous proteins and clarifying their limitations in *R. palustris* advances the efforts to engineer this robust microbe.

## Materials and Methods

### Strain Growth Conditions

*Rhodopseudomonas palustris* (Molisch) van Niel BAA-98, strain designation CGA009 (*R. palustris*), was obtained from American Type Culture Collection. NEB® 10-beta competent *Escherichia coli (E. coli* DH10β) was used for plasmid construction. All strains used in this study are described in Supplementary Table I, and were stored at −80°C, *R. palustris* strains were stored with a final concentration of 20% glycerol while *E. coli* strains were stored at a 15% final glycerol concentration. Before growth in liquid media, *E. coli* and *R. palustris* strains were grown on solid LB media (Miller, AMRESCO) and 112 Van Niel’s (VN) media (ATCC) (1% yeast extract, 0.1% K_2_HPO_4_, 0.05% MgSO_4_, pH 7.1) plates respectively with the appropriate antibiotic (listed below). All strains were grown in 4 mL of their respective media in 14 mL BD Falcon™ round-bottom polystyrene tubes at 275 rpm in the dark in ambient air at 30°C. *E. coli* cultures containing plasmids with ampicillin (amp), gentamicin (gent), or kanamycin (kan) selection markers were grown with 100μg/mL amp, 10 μg/mL gent, or 30 μg/mL kan, respectively. *R. palustris* cultures were grown in 112 Van Niel’s (VN) media (ATCC) or photosynthetic media (PM) (Brown et al. 2020) supplemented with 20 mM NaC_2_H_3_O_2_, 10 mM NaHCO_3_, and 15.2 mM (NH_4_)_2_SO_4_ where specified. *R. palustris* cultures containing plasmids with the corresponding selection markers were grown with 25, 50, or 100 μg/mL amp, 300 μg/mL gent, or 300 μg/mL kan.

### Strain Construction

Oligonucleotides were purchased from Eurofins Genomics or Integrative DNA Technologies™. All plasmids used in this work are listed in Supplementary Table I. The sequences of the genetic parts are listed in Supplementary Table II. PCR was conducted using Phusion Hot Start II DNA Polymerase (Thermo Scientific™). On condition that the PCR product was the desired size as determined through gel electrophoresis with 1X TAE buffer 1% agarose gels, the PCR reaction was purified via Monarch® DNA Gel Extraction Kit (New England Biolabs® Inc.). Plasmids were constructed via the Hot Fusion Assembly Method (Fu et al., 2014) and used to transform *E. coli* DH10β. *E. coli* was grown overnight, diluted to 1/40 OD_600_ in fresh LB media, grown until mid-exponential phase, and washed at room temperature according published literature (Tu et al., 2016). Washed cells were then transformed by the Hot Fusion product through electroporation. Electroporated cells were incubated at 30°C for 1.5 hours in LB media without antibiotic, then plated onto LB plates supplemented with appropriate antibiotic. Plates were incubated overnight at 30°C. Colonies were then selected and cultures were grown in LB media overnight with the appropriate antibiotic. Cultures were stored as 15%(v/v) glycerol stock at −80°C. Plasmids were harvested from the cultures using the PureLink™ Quick Plasmid Miniprep Kit (Invitrogen™). After extraction of the assembled plasmids, junctions in the plasmids were confirmed by submission of PCR products to Eurofins Genomics for DNA sequencing. Plasmids were then used to transform *R. palustris* through the same method with the following exceptions. *R. palustris* was diluted to 0.2 OD_660_ and grown overnight. After electroporation, *R. palustris* was incubated in VN media without antibiotic overnight, then plated onto VN plates supplemented with appropriate antibiotic. Plates were incubated approximately 5-7 days after transformation until colonies emerged. These colonies were then streaked onto fresh VN plates supplemented with appropriate antibiotic and allowed to grow approximately five days. Liquid cultures were grown from these plates in VN media with appropriate antibiotic. *R. palustris* cultures were stored at −80°C at a final concentration of 20%(v/v) glycerol. The template for Colony PCR was cells suspended in water and lysed at 100°C for 25 minutes. PCR products of sequences critical for performance were confirmed by submission of PCR products to Eurofins Genomics for DNA sequencing.

Sucrose counter selection was used to remove the chloramphenicol acetyltransferase gene from *R. palustris*. Briefly, a suicide plasmid was constructed using the p15A origin of replication. The plasmid (pΔcat) contained two adjacent 1300-1500 bp homology arms to allow for integration into *R. palustris*’ genome and are described in Supplementary Table III. This plasmid also contained a gentamicin resistance gene to provide selection pressure and a *sacB* gene to confer sucrose lethality. This plasmid was used to transform *R. palustris* following the previously described method and plated onto VN/gentamicin plates (300 μg/mL gent). Gentamicin-resistant colonies were then picked and grown in 4 mL PM with 10 mM sodium succinate without gentamycin. Cultures were grown at 30°C and 275 rpm for two days to allow for recombination. Cells were then removed from cultures through centrifugation and resuspended in 200 mL PM media. Serial dilutions of 1/1,000 and 1/10,000 were then performed on these cultures and dilutions were plated on PM media plates supplemented with 10% sucrose. Plates were incubated for 6 days to allow colonies to emerge. Colonies were then tested on duplicate grid plates. This was done by transferring a colony to a PM plate supplemented with 215 μg/mL gentamicin and simultaneously transferring to a plain PM plate to determine a loss of vector-mediated gentamicin-resistance. Colonies that did not grow on PM plates supplemented with gentamicin, but did grow on plain PM media plates were then selected and transferred into culture tubes containing 5 mL PM. Cultures were then incubated at 30°C for 6 days until a purple color appeared. Colony PCR with the segregation primers listed in Supplementary Table IV was performed and sequencing via submission to Eurofins Genomics was used to verify the deletion strain.

Mutations to *R. palustris*’ endogenous plasmid was accomplished through double homologous recombination. Briefly, a suicide plasmid was constructed from the expression cassette flanked by two 800-1000 bp homology arms and the p15A origin of replication (ori). All homology arm sequences are described in Supplementary Table III. Colony PCR was used to validate that the ori does not replicate in *R. palustris*. The homology arms were amplified from *R. palustris* gDNA which was obtained using Monarch® Genomic DNA Purification Kit (New England Biolabs® Inc.). A gentamicin selection marker was included in the expression cassette to force segregation into the endogenous plasmid. Electroporation of the suicide plasmid and culturing of the subsequent colonies was conducted following the previously described methods. Colony PCR with the segregation primers listed in Supplementary Table IV was performed and sequencing via submission to Eurofins Genomics was used to verify the incorporation of the expression cassette.

### Antibiotic Tolerance Determination

Wild type *R. palustris* cultures were grown in 50 mL of PM media supplemented with 20 mM NaC_2_H_3_O_2_, 10 mM NaHCO_3_, 15.2 mM (NH_4_)_2_SO_4_, in 250 mL Erlenmeyer flasks to stationary phase. Cultures were then diluted to 0.2 OD_660_ into 70 mL of fresh PM media supplemented with 20 mM NaC_2_H_3_O_2_, 10 mM NaHCO_3_, 15.2 mM (NH_4_)_2_SO_4_, and one of the following antibiotics; 50, 100μg/mL amp, 34 μg/mL cm, 300 ug/mL gent, 200, 300 μg/mL kan, or 300 μg/mL spec. The cultures were then transferred into glass 85mL tubes, and placed into a Multi-Cultivator 1000-OD (Photon Systems Instruments). Theses cultures were incubated at 30°C, in the dark, and bubbled with ambient air. The cultures’ absorbance (680 nm) was measured by the Multi-Cultivator 1000-OD every two hours until it did not change for at least 3 measurements.

### Fluorescence Fold Change Measurement

*R. palustris* strains were grown to stationary phase in PM media supplemented with 20 mM NaC_2_H_3_O_2_, 10 mM NaHCO_3_, 15.2 mM (NH_4_)_2_SO_4_ and the appropriate antibiotic in 250 mL Erlenmeyer flasks before dilution to 0.2 OD_660_ in 500μL of fresh media and the appropriate antibiotic concentration, in triplicate. The cultures were loaded into a Greiner CELLSTAR® bio-one sterile 48 well culture plate and covered with a Breathe-Easy® Gas Permeable Sealing Membrane (Diversified Biotech). Cultures were then incubated in the dark at 30°C in ambient air for 72 hours at 275 rpm. Afterwards, 200 μL of each culture was pipetted into a Greiner bio-one 96 well polystyrene flat-bottomed μCLEAR® black microplate before fluorescence of reporter proteins and absorbance of the cultures were measured. Three wells were also loaded with PM media to act as blanks. Plate was then loaded into a Molecular Devices SpectraMax® i3x microplate reader. Excitation-Emission wavelengths for mRFP, eYFP, and GFPuv were respectively: 583-608 nm, 485-528 nm, and 395-509 nm. Absorbance of cultures at 660nm was also measured. The following equation was used to calculate each strain’s relative fluorescence.

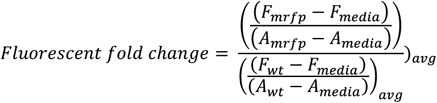

### Flow Cytometry

*R. palustris* strains were grown according the Fluorescent Fold Change Measurement section. Cultures were then resuspended in 150 μL 0.85% NaCl at 0.001 OD_660_ and loaded into a Fisherbrand clear, polystyrene, 350 μL, flat-bottom 96 well plate. The cultures were analyzed by a Beckman Coulter CytoFLEX LX flow cytometer. mRFP was excited by a 561 nm yellow/green laser and the emission was collected using a 610/20 nm bandpass filter. 10,000 bacterial events were collected per sample.

### Terminator Efficiency

The *R. palustris* strains were grown and their fluorescence and absorbance determined according the Fluorescent Fold Change Measurement section. The percent reduction in read through was calculated by the following equation.

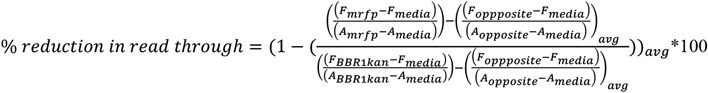

### Construction of the RBS Library and 5’ UTR Replacement

To construct the Ribosome Binding (RBS) library, *mrfp*’s six base pair RBS was subjected to saturation mutagenesis using degenerate oligonucleotides for PCR. The Blunt End Ligation Method was used to construct the plasmid (Georgescu et al., 2003) with T4 DNA Ligase and T4 Polynucleotide Kinase (New England Biolabs). The sequence of the RBS of selected colonies was determined by submission of colony PCR products to Eurofins Genomics for DNA sequencing (Supplementary Table V). The De Novo DNA RBS calculator was used to predict the translation initiation rate of the *mrfp* 5’ UTRs (Cetnar, Salis, 2021; Reis, Salis, 2020, Espah Borujeni, Salis, 2016; Espah Borujeni et al., 2014; Espah Borujeni et al., 2017; Salis et al., 2009). The De Novo DNA RBS calculator for controlling translation initiation rates was then used to design the 5’ UTR sequences for the eYFP plasmids. The 5’ UTRs were exchanged using Blunt End Ligation method. All 5’ UTR sequences are listed in Supplementary Table II.

### RNA Extraction, gDNA Removal, and Reverse Transcription

RNA was extracted from *R. palustris* cultures at mid-exponential phase in duplicate according to TRIzol® Reagent (Life Technologies™) protocol, with the following modifications. After the addition of chloroform required for phase separation, cells were incubated at room temperature for 5 minutes. After incubation of samples with isopropanol to precipitate the RNA, cells were centrifuged for 30 minutes instead of 10 minutes at 12000g and 4°C. Ice cold 75% ethanol was used to wash RNA pellets. After the RNA wash, samples were centrifuged for 10 minutes at 7500g and 4°C.

RNA extracts were treated with TURBO™ DNase (Invitrogen™) to remove any gDNA that may have also been extracted. Afterwards, PCR of the RNA with the 16SrRNA primers (Supplementary Table VI) and gel electrophoresis with a 2% agarose gel was used to confirm the absence of any gDNA. A bleach gel was then used to verify the integrity and lack of degradation of RNA after extraction, as described in literature (Aranda et al., 2012). Briefly, 1 μg total RNA was run in a 1X TAE 1% agarose gel with 0.5% (v/v) bleach. Gel electrophoresis was conducted and gels were imaged to verify RNA integrity. Wells that contained two equivalent bands after DNase treatment were considered to not be degraded. RNA samples that were not degraded were converted to cDNA in 20 μL reactions using a High-Capacity cDNA Reverse Transcription Kit (Applied Biosystems™).

### RT-qPCR

All oligonucleotides used for RT-qPCR are outlined in Supplementary Table VI. Primer concentrations for RT-qPCR reactions were initially evaluated by performing 25μL PCR reactions using Go Taq® Master Mix (Promega Corporation) with approximately 100μg of gDNA, for primer concentrations ranging from 350nM to 50nM. A second set of reactions containing no gDNA was also prepared. Thermo cycler settings for this reaction were 95°C for 2 minutes, 40 cycles of (95°C for 45 seconds, 60°C for 45 seconds, 72 for X seconds – based on amplicon length), and then 72°C for 5 minutes. Gel electrophoresis was then performed on PCR products using 1X TAE 2% agarose gels. Primer concentrations that produced a band in reactions containing gDNA but no band in the second reaction set were selected for future evaluation.

Primer efficiency was determined by preparing qPCR reactions with PowerUp SYBR® Green Master Mix (Life Technologies™), with a 5x dilution series of cDNA (500ng/μL - 0.0061ng/μL). The primer concentration for each target gene for qPCR reactions is also listed in Supplementary Table VI. qPCR reactions were run in an Eppendorf Mastercycler Realplex. Thermocycler settings for qPCR reactions were 50°C for 2 minutes 95°C for 2 minutes, 40 cycles of (95°C for 15 seconds, 60°C for 1 minute), with a melting curve step (60°C for 1 minute, rising at a rate of 1.5°C for 20 minutes, staying at 95°C for 15 seconds) to ensure a single amplicon. Non-template controls (NTC) in triplicate were included for each reaction to ensure a lack of unintended PCR products for each target gene. The Eppendorf Mastercycler Realplex automatic baseline calculator was used to calculate the baseline for samples. Cycle threshold (CT) values were then obtained from the Eppendorf Mastercycler Realplex. Primer efficiency for each primer set reaction was determined by plotting the Log10 of cDNA copies vs the CT value for each set of reactions. Linear regression was used to ensure linearity of this plot. The slope of this graph determined the primer efficiency of the reaction. The ThermoFisher Scientific qPCR calculator was used to determine primer efficiency for each reaction set. Using this method, all primers were ensured to have a primer efficiency between 90%-110%., reported in Supplementary Table VI.

qPCR reactions for two biological replicates and two technical replicates were then run in the Eppendorf Mastercycler Realplex using the same settings as for primer efficiency determination. The concentrations of primers and cDNA are listed in Supplementary Table VI. NTC controls in triplicate were also run. The relative mRNA concentration was calculated per the following equation, GOI is the gene of interest and HK is the housekeeping gene.

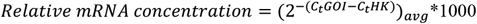

### Plasmid Copy Number

qPCR with gDNA was used to determine the absolute plasmid copy number ratio of both native and non-native plasmids relative to the chromosome as described previously (DeLorenzo et al., 2018). gDNA was extracted from *R. palustris* cultures at mid-exponential phase in duplicate using the Monarch® Genomic DNA Purification Kit (New England Biolabs® Inc.). Amplified PCR product using primers in Supplementary Table VI was diluted to 10^9^ copies of amplicon/μL per the following equation.

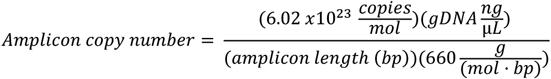

An external standard curve was constructed using a 5x dilution series starting with 10^7^ copies/uL in triplicate. The primer efficiency was determined as outlined in the previous section.

qPCR reactions for two biological replicates and two technical replicates were then run in the Eppendorf Mastercycler Realplex using the same settings as for primer efficiency determination. The concentrations of primers and gDNA are listed in Supplementary Table VI. NTC controls in triplicate were also run. The standard curve was used to determine the number of copies of each amplicon. The plasmid copy number relative to the chromosome was then determined.

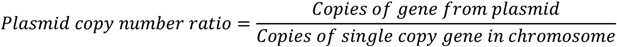

### Statistical Methods

All statistical analyses (Student’s two-tailed t-test with unequal variances) was performed using Microsoft Excel. p-value < 0.05 was considered to be significant. All experiments were conducted in biological triplicate, with exception for RT-qPCR and plasmid copy number, which was performed in biological duplicate and technical duplicate.

## Results

### Sensitivity to antibiotics used for selection

Engineering *R. palustris* to explore its unique biochemical abilities has required very high concentrations of the antibiotic used for selection (Braatsch et al., 2006;Rey et al., 2006;Huang et al., 2010;Pechter et al., 2016;Doud et al., 2017); yet, characterization of *R. palustris*’ sensitivity to the antibiotics commonly used for genetic engineering was not found in literature searches. To address this problem, either ampicillin, kanamycin sulfate, gentamicin sulfate, or spectinomycin sulfate was added to wild type *R. palustris*’ media and the absorbance (680 nm) was recorded every two hours. The maximum change in absorbance and the time to achieve that maximum change was then determined (Figure 1a). *R. palustris* was most sensitive to ampicillin, with both 50 μg/mL and 100 μg/mL eliminating the bacterium within ten hours. Kanamycin selection has been commonly used when engineering *R. palutris*; yet, it took over 30 hours for the growth (average change in absorbance of 0.1) to completely stop after the addition of either 200 μg/mL or 300 μg/mL. The cultures without any antibiotics reached stationary phase in a similar amount of time. It was also more than 30 hours for the absorbance of the cultures with 300 μg/mL spectinomycin to stop changing; although, the average maximum change was only 0.03. Gentamicin has also been used for selection to engineer *R. palustris*. Adding 300 μg/mL to the *R. palustris* cultures arrested growth in an average of 13 hours while the absorbance increased by just 0.05.

**Figure 1.**
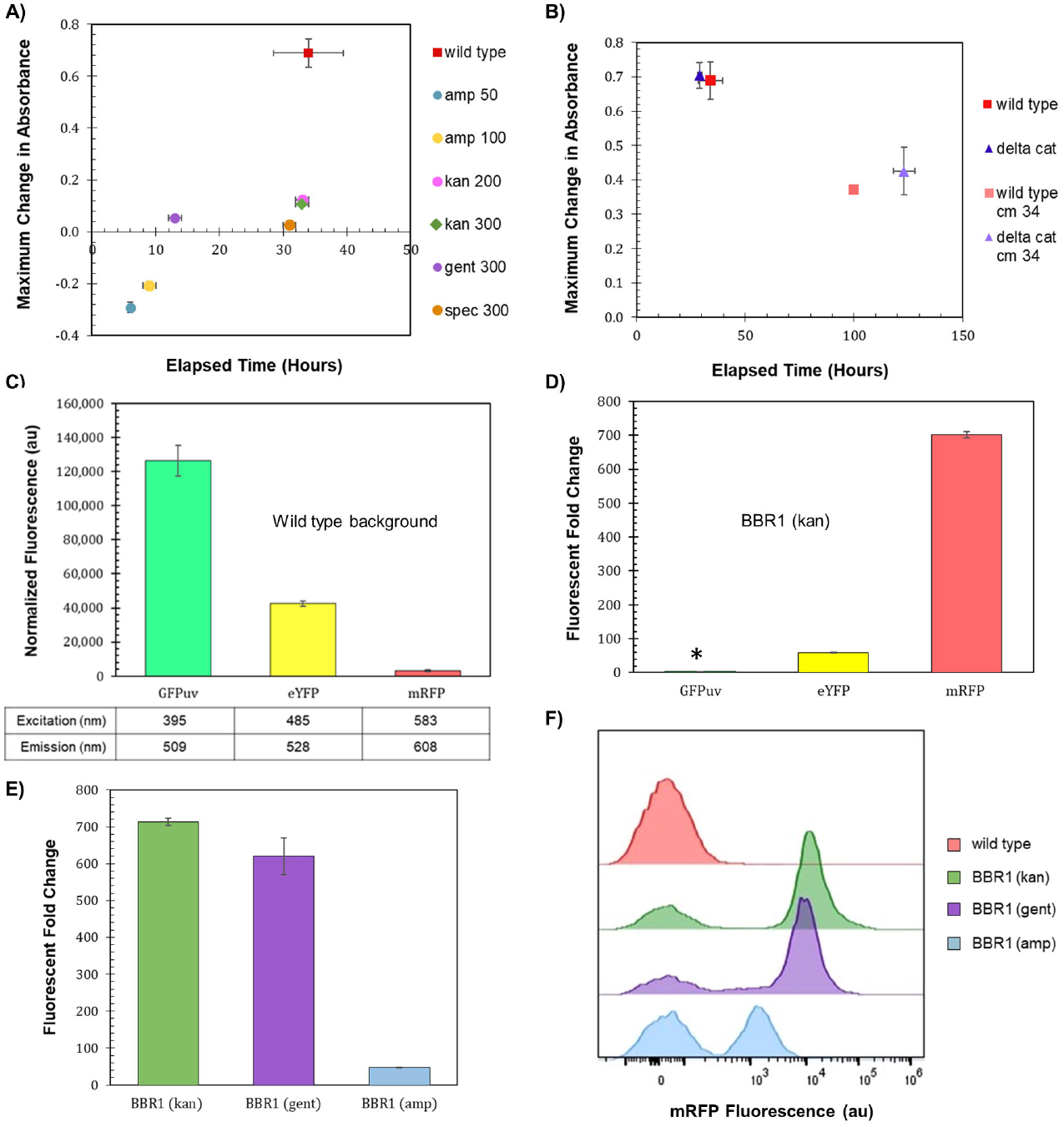
Selection marker and fluorescent reporter characterization. **A)** Wild type *R. palustris*’ change in absorbance in response to the addition of antibiotics (commonly used in synthetic biology) to the media and growth recorded over time. Wild type is *R. palustris* without any antibiotics added to the media. Amp 50 and Amp 100 are 50 μg/mL and 100 μg/mL ampicillin respectively. Kan 200 and Kan 300 are 200 μg/mL and 300 μg/mL kanamycin sulfate respectively. Gent 300 and Spec 300 are 300 μg/mL gentamicin sulfate and 300 μg/mL spectinomycin sulfate respectively. **B)** Change in absorbance for wild type *R. palustris* and the *R. palustris* Δ*cat* strain in response to the addition of 34 μg/mL chloramphenicol added to the media and growth recorded over time. **C)** Average fluorescence of wild type *R. palustris* normalized to absorbance (660 nm), in triplicate, at the emission and excitation wavelengths for GFPuv, eYFP, and mRFP (Materials and Methods). Error bars indicate one standard deviation. **D)** Fluorescent fold change average of *R. palustris* BBR1 kan GFPuv, BBR1 kan eYFP, and BBR1 kan mRFP strains grown in triplicate. 300 μg/mL kanamycin sulfate was added to the media for the mutant strains but not wild type *R. palustris* (Materials and Methods). * indicates statistically significant expression of 3-fold for the BBR1 kan GFPuv strain. Error bars indicate one standard deviation. **E)** Fluorescent fold change average of *R. palustris* BBR1 kan mRFP, BBR1 gent mRFP, and BBR1 amp mRFP strains grown in triplicate. The antibiotic concentrations added to the media of the mutant strains but not wild type *R. palustris* were 300 μg/mL kanamycin sulfate, 300 μg/mL gentamicin sulfate, and 50 μg/mL ampicillin respectively (Materials and Methods). Error bars indicate one standard deviation. **F)** Representative flow cytometry results of mRFP fluorescence from *R. palustris* wild type, BBR1 kan mRFP, BBR1 gent mRFP, and BBR1 amp mRFP strains grown in triplicate. The growth procedure, media, and antibiotics are the same as for the fluorescent fold change test in Figure 1e (Materials and Methods).

A chloramphenicol acetyltransferase (CAT), which inactivates chloramphenicol, is encoded in *R. palustris*’ chromosome. Furthermore, *cat* is expressed in both aerobic and anaerobic conditions (Supplementary Figure 1) as verified by PCR of the cDNA (Materials and Methods). In an effort make *R. palustris* sensitive to chloramphenicol, *cat* was removed from the chromosome using sucrose counterselection (Materials and Methods). Wild type and Δ*cat R. palustris*’ sensitivity to chloramphenicol was then tested, following the same protocol used to test *R. palustris*’ sensitivity to the other antibiotics. Both strains’ growth was hampered by the antibiotic, but still reached a maximum absorbance that was 50% of the same strains grown without chloramphenicol (Figure 1b).

### Fluorescent reporters for parts characterization

Fluorescent reporters are a fundamental synthetic biology tool for characterizing genetic parts in vivo (Delvigne et al., 2015). The background fluorescence produced by photosynthetic microbes like *R. palustris* can complicate the use of such an important tool. The gene for green fluorescent protein (GFP) has been expressed in *R. palustris* CGA009 (Doud et al., 2017) as well as another *R. palustris* strain, GJ-22 (Zhai et al., 2019), but the use of other fluorescent proteins has not been reported. To determine the fluorescent reporter best suited for work in *R. palustris*, the fluorescence of wild type *R. palustris* was determined (Materials and Methods) for the excitation and emission wavelengths of GFPuv, enhanced yellow fluorescent protein (eYFP), and monomeric red fluorescent protein (mRFP) (Figure 1c). Interestingly, the pink pigmented bacterium’s background fluorescence was lowest for mRFP. Next, *R. palustris* transformed by a plasmid using a kanamycin selection marker, the pBBR1 origin of replication (Kovach et al., 1995), and either *gfpuv* (Lee et al., 2011), *eyfp* (Knoot et al., 2019), or *mrfp* (Bi et al., 2013) expressed from the *lac* promoter was tested following the protocol for measuring the background fluorescence of the wild type strain. A pBBR1-based plasmid was chosen since they have been used for heterologous gene expression previously in *R. palustris* (Braatsch et al., 2006;Huang et al., 2010;Heiniger and Harwood, 2015). The strain expressing *gfpuv* produced a statistically significant normalized fluorescence as compared to the wild type strain’s normalized background fluorescence (student’s two-tail t-test, p < 0.05) of only three-fold while the fluorescent fold change for the *eyfp*-expressing strain was 55 (Figure 1d). To verify expression of *gfpuv*, RNA was extracted from cultures of the *gfpuv*-expressing strain at mid-exponential phase, in addition to wild type *R. palustris* cultures. PCR of the subsequent cDNA shows expression just from the pBBR1 kan GFPuv cultures. (Supplementary Figure 1). The *mrfp*-expressing strain produced more than 700-fold higher fluorescence than the wild type background and was thus selected for future work.

### Selection marker testing

300 μg/mL kanamycin was sufficient for the *mrfp*-expressing *R. palustris* to produce a fluorescent fold change of 700. Both ampicillin and gentamicin allowed a smaller positive change in absorbance in a shorter time as compared to kanamycin when testing the microbe’s sensitivity to the antibiotics. Therefore, the kanamycin selection marker (kan) in the pBBR1 plasmid was replaced with an ampicillin resistance gene (amp) or a gentamicin resistance gene (gent) and used to transform *R. palustris*. The fluorescent fold change from the gent strain was statistically similar to the kan strain (student’s two-tail t-test, p > 0.05), while the amp strain produced just a 48-fold change in fluorescence (Figure 1e). Flow cytometry was then employed to investigate the differences in the mRFP-producing strains as compared to the wild type *R. palustris*, all in triplicate (Materials and Methods). All cultures included some cells with just the background fluorescence of the wild type strain. Figure 1f is representative of the results. The averages and standard deviations are presented in Figure 6f. More than 50% of the cells measured produced just the background fluorescence for the amp strain with 50 μg/mL amp. The pBBR1 amp strain was also tested with 25 and 100 μg/mL ampicillin. The average fluorescent fold change was statistically lower (student’s two-tail t-test, p < 0.05) and the variation was larger when 25 μg/mL ampicillin was used (Supplementary Figure 2). When the ampicillin concentration was increased to 100 μg/mL, all of the cells just produced the wild type background fluorescence (Supplementary Figure 3). For the pBBR1 kan and pBBR1 gent strains there were fewer cells with just the background fluorescence, with an average just under 20%. Either a kanamycin or gentamicin selection marker was included in all of the remaining plasmids built for this study.

### Origins of replication

Plasmids with the pBBR1 origin of replication (ori) have been commonly used for heterologous gene expression in *R. palustris* (Braatsch et al., 2006;Huang et al., 2010;Heiniger and Harwood, 2015). Yet, studies that determine how well pBBR1 plasmids are maintained by *R. palustris* were not found in literature. Fluorescent fold change, flow cytometry, and the relative quantification of plasmid copy number were utilized to access pBBR1’s stability under kanamycin or gentamicin selection (Figure 2). Plasmid copy number was of interest because it directly affects the number of proteins produced from the genes encoded on the plasmid, as well the stable distribution of the vector to daughter cells during growth (Jahn et al., 2016). The same tests were also conducted after exchanging the pBBR1 with the RSF1010 replicon, another origin of replication with a broad host range (Taton et al., 2014;Knoot et al., 2019).

**Figure 2.**
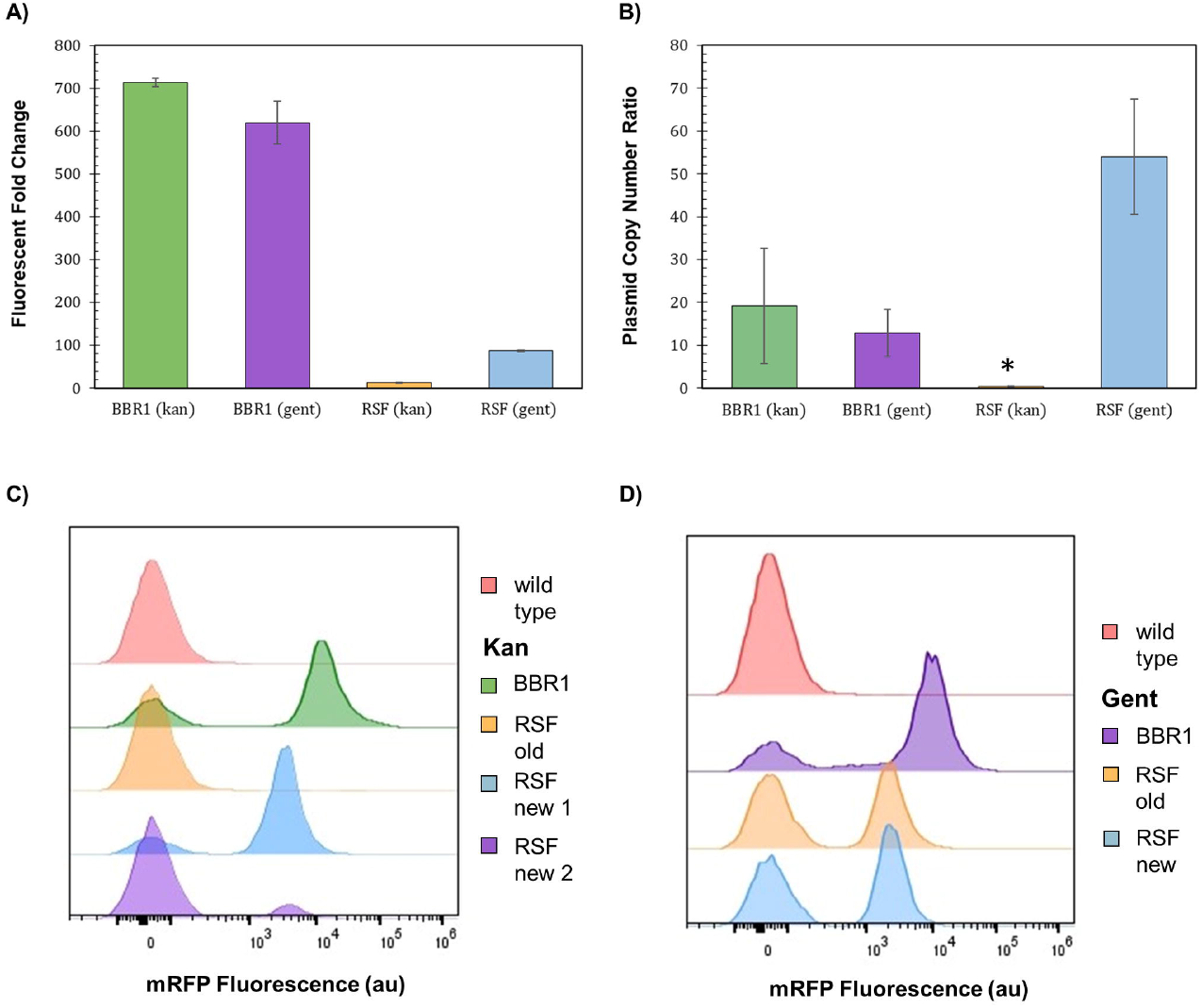
Origin of replication characterization with different selection markers. **A)** Fluorescent fold change average of *R. palustris* BBR1 kan mRFP, BBR1 gent mRFP, RSF1010 kan mRFP, and RSF1010 gent mRFP strains grown in triplicate. The antibiotic concentrations added to the media of the mutant strains but not wild type *R. palustris* were 300 μg/mL kanamycin sulfate and 300 μg/mL gentamicin sulfate respectively (Materials and Methods). Error bars indicate one standard deviation. **B)** Average plasmid copy number for the *R. palustris* BBR1 kan mRFP, BBR1 gent mRFP, RSF1010 kan mRFP, and RSF1010 gent mRFP strains grown in duplicate. The antibiotic concentrations added to the media of the mutant strains were 300 μg/mL kanamycin sulfate and 300 μg/mL gentamicin sulfate respectively. Copies of the kanamycin and gentamycin selection marker gene were compared to the single copy *dxs* gene in the chromosome by qPCR and an external standard curve. * indicates a statistically significant plasmid copy number of 12.1-fold for the RSF1010 kan mRFP strain. (Materials and Methods). Error bars indicate one standard deviation. **C)** Representative flow cytometry results of mRFP fluorescence from *R. palustris* wild type, BBR1 kan mRFP, and RSF1010 kan strains grown in triplicate. The old RSF1010 kan cultures had been streaked from frozen stock two weeks earlier. The new RSF1010 kan cultures were single colonies from a new transformation. The growth procedure, media, and antibiotics are the same as for the fluorescent fold change test in Figure 2a (Materials and Methods). **D)** Representative flow cytometry results of mRFP fluorescence from *R. palustris* wild type, BBR1 gent mRFP, and RSF1010 gent strains grown in triplicate. The old RSF1010 gent cultures had been streaked from frozen stock two weeks earlier. The new RSF1010 gent cultures were single colonies from a new transformation. The growth procedure, media, and antibiotics are the same as for the fluorescent fold change test in Figure 2a (Materials and Methods).

As determined when testing the selection markers, the fluorescent fold change for pBBR1 plasmids expressing *mrfp* using kan or gent selection were statistically similar, averaging about 650-fold (Figure 2a). There was a significant difference in the fluorescent fold change for the RSF1010 plasmids with the two selection markers, 87-fold for gent and only 12-fold for kan. Flow cytometry of the cultures in triplicate indicate that the RSF1010 plasmid with kan selection is not stably maintained in *R. palustris*. Figure 2c is representative of the results. The averages and standard deviations are presented in Figure 6f. Only cells that produced background fluorescence were found in a RSF1010 kan culture that had been streaked from frozen stock two weeks earlier. In addition, one culture from a newly transformed RSF1010 colony was predominantly fluorescent cells, 82%, while the number of fluorescent cells in a culture grown from a second colony from the same transformation was just 12%. This variation in cultures harboring the RSF1010 plasmid was not seen when gentamicin was used for selection (Figure 2d). The average number of cells from the RSF1010 gent cultures with just background fluorescence was higher than for the pBBR1 gent cultures, 50% for RSF1010 gent and 20% for pBBR1 gent, but unlike the RSF1010 kan cultures, that percentage was not dependent on the age of the cultures.

The copy number of the pBBR1 and RSF1010 plasmids, with kan and gent selection markers, was determined relative to *R. palustris*’ number of chromosomes by qPCR with the cultures’ gDNA and standard curves derived from a dilution series of each PCR product (Materials and Methods) (Lee et al., 2006;DeLorenzo et al., 2018). A section of each selection marker and of a single copy gene in the chromosome were amplified for the comparison. Similar to the fluorescent fold change and flow cytometry results, the relative plasmid number was equivalent for the pBBR1 plasmids. There were approximately 15 copies per chromosome for both pBBR1 plasmids (Figure 2b). Also similar to the results derived from mRFP fluorescence, there was a significant difference in the relative plasmid copy number for the RSF1010 plasmids, less than one copy per chromosome for the RSF1010 kan cultures and averaging over 50 copies per chromosome for the RSF1010 gent cultures.

### Aerobic versus phototrophic/anaerobic gene expression

*R. palustris* is photosynthetic in anaerobic conditions, providing the organism energy from light in addition to the energy it can obtain from organic compounds (Larimer et al., 2004). To take advantage of this important source of energy, synthetic biology tools for heterologous gene expression also need to be characterized during phototrophic growth. The most commonly used fluorescent reporters, developed from the jellyfish *Aequorea victoria*’s GFP, require oxygen to synthesize their chromophores (Drepper et al., 2007). In addition, the oxygen-independent flavin-binding fluorescent proteins are not bright enough to be clearly discernable above *R. palustris*’ background fluorescence at their excitation and emission wavelengths (Mukherjee et al., 2013). Therefore, RNA and gDNA was extracted from the *R. palustris* pBBR1 kan mRFP strain grown aerobically and anaerobically (100 μE white light) for RT-qPCR (following the MIQE guidleines (Bustin et al., 2009)) and plasmid copy number tests (Materials and Methods). In addition, RNA was extracted from the pBBR1 gent mRFP, RSF1010 kan mRFP, and RSF1010 gent mRFP strains, and wild type *R. palustris* controls, all grown to mid-exponential phase.

Large variation in both *mrfp* expression relative to the reference gene and plasmid copy number was found for the biological replicates of the pBBR1 kan mRFP strain (Figure 3). The average relative *mrfp* expression for the pBBR1 kan cultures grown aerobically was minimal when compared to expression from the anaerobically grown cultures (Figure 3a). The aerobic relative *mrfp* expression from the pBBR1 kan mRFP strain was also compared to the other strains grown aerobically, (pBBR1 gent, RSF1010 kan and RSF1010 gent) (Supplementary Figure 4). The *mrfp* expression of the aerobically grown strains was similar to the fluorescent fold change results (Figure 2a); the cultures for the two pBBR1 ori strains had higher average relative *mrfp* expression than the RSF1010 gent culture, and the RSF1010 kan culture had no detectable *mrfp* expression. The plasmid copy number more than doubled to 50 copies per chromosome with a huge variation between replicates, when comparing the pBBR1 kan mRFP cultures grown anaerobically to those grown aerobically (Figure 3b).

**Figure 3.**
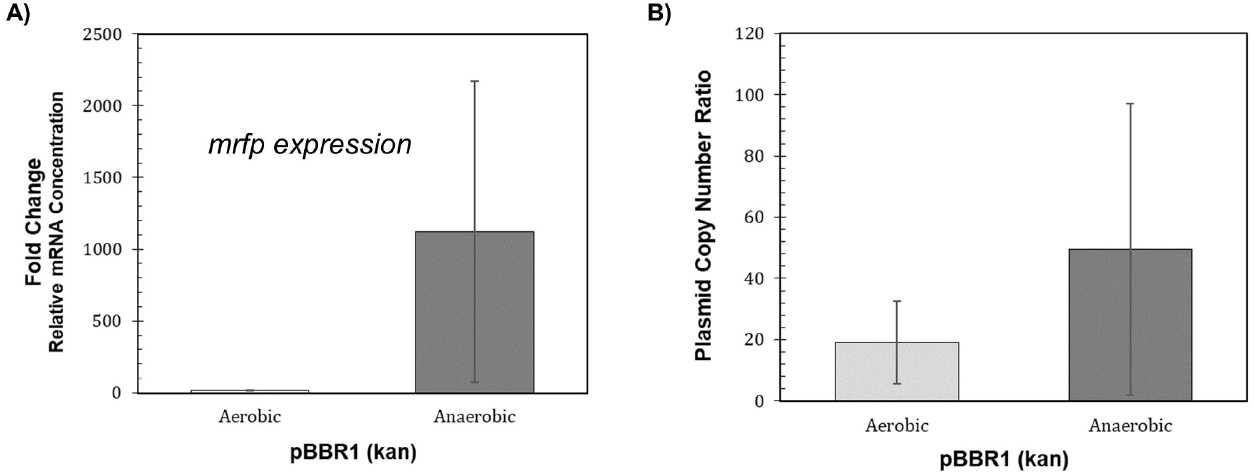
Performance of the *R. palustris* BBR1 kan mRFP strain during phototrophic growth. **A)** Relative mRNA concentration (*mrfp* relative to *16rRNA*) for the *R. palustris* BBR1 kan strain when grown aerobically in the dark and anaerobically in the light (100 μE white light) (Materials and Methods). Two biological and two technical replicates were averaged for each strain. Error bars indicate one standard deviation. **B)** Average plasmid copy number for the *R. palustris* BBR1 kan mRFP strain grown aerobically in the dark and anaerobically in the light (100 μE white light) in duplicate. Copies of the kanamycin selection marker gene was compared to the single copy *dxs* gene in the chromosome by qPCR and an external standard curve (Materials and Methods). Error bars indicate one standard deviation.

### Genetic context and transcriptional terminators

Transcription does not necessarily stop at the end of a gene, but can continue on to other genes on the same DNA strand (Chen et al., 2013;Kelly et al., 2019). The fluorescent reporter gene on the original pBBR1 plasmids was preceded on the same DNA strand by the antibiotic resistance gene, which was not true for the same fluorescent reporter gene on the RSF1010 plasmids. Two transcriptional terminators, tonB and rrnC, were tested for their ability to reduce the transcriptional read-through from the antibiotic resistance gene, and both were compared to the strain with the *P_Lac_-mrfp* expression cassette on the opposite strand of DNA in the pBBR1 kan plasmid (Materials and Methods). Both terminators significantly reduced transcriptional read-through in *E. coli*; although, the rrnC terminator was determined to be 25 times stronger than tonB (Chen et al., 2013).

Moving the *P_Lac_-mrfp* expression cassette to on the opposite strand of DNA in the pBBR1 kan plasmid, reduced the fluorescent fold change more than eight times to 82 (Figure 4), which is similar to the fluorescent fold change from the RSF1010 gent plasmid, at 87 (Figure 2a). Inserting the tonB terminator between *kan* and *mrfp* resulted in a three-fold decrease in fluorescent fold change while the rrnC terminator produced more than a six-fold reduction. The strain with the expression cassette on the opposite strand was used to calculate how much of the mRFP fluorescence was due to transcription of the antibiotic resistance gene (read-through). The ratio of normalized mRFP fluorescence for the tonB strain and the rrnC strain to the pBBR1 kan strain without a terminator was used to calculate the percent reduction in read-through (Materials and Methods). The rrnC terminator was stronger than the tonB terminator, reducing read-through by 96% versus 75%.

**Figure 4.**
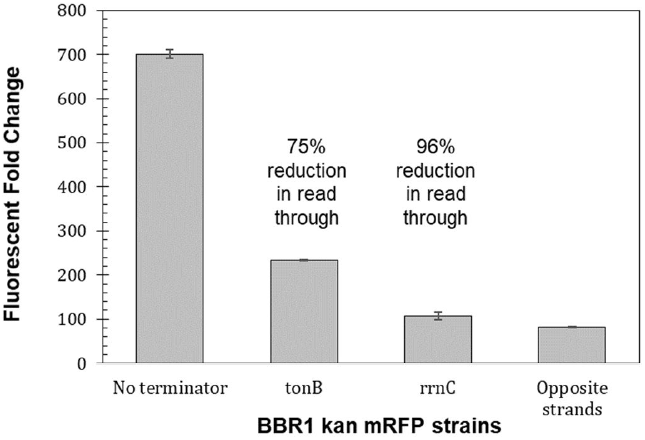
Transcriptional terminators reduce read-through from the preceding transcription unit. Fluorescent fold change average of *R. palustris* BBR1 kan mRFP (no terminator), BBR1 kan mRFP tonB, BBR1 kan mRFP rrnC, and BBR1 kan mRFP (opposite strand) strains grown in triplicate (Materials and Methods). Error bars indicate one standard deviation.

### Predicting changes in protein production based on the 5’ UTR

The 5’ untranslated region (UTR) of mRNA significantly impacts protein production through its participation in transcription and the stability of the transcript, as well as translation (Balzer Le et al., 2020). To investigate the dynamic range of that could be achieved and whether the level of fluorescent protein production could be predicted, the six bases of *mrfp*’s ribosome site (RBS) were randomized on the pBBR1 kan plasmid (Materials and Methods). A scan of 450 colonies yielded a dynamic range of 562-fold (Figure 5a). 32 colonies from across the range of normalized fluorescence, plus the original pBBR1 kan mRFP strain, were then tested per the Fluorescent Fold Change Measurement protocol, which produced nearly a 1,300-fold dynamic range (Figure 5b). The sequence of the RBS was determined for the 32 colonies and used to generate a predicted translation initiation rate (TIR) with the DeNovo DNA RBS calculator (Salis et al., 2009;Espah Borujeni et al., 2014;Espah Borujeni and Salis, 2016;Espah Borujeni et al., 2017;Reis and Salis, 2020;Cetnar and Salis, 2021). The predicted TIR versus the fluorescent fold change was then plotted for each strain (Figure 5c). Since there can be variability in protein production even within biological replicates, the graph was divided into nine even sections (3 × 3) to help analyze the data. In general, the 5’ UTRs with the lowest TIRs resulted in strains with the lowest fluorescent fold change. When the TIR of the 5’ UTR was in the middle range, the fluorescent fold change spanned the entire dynamic range. A high TIR resulted in two colonies that bordered on the high fluorescent fold change range and two in the middle range.

**Figure 5.**
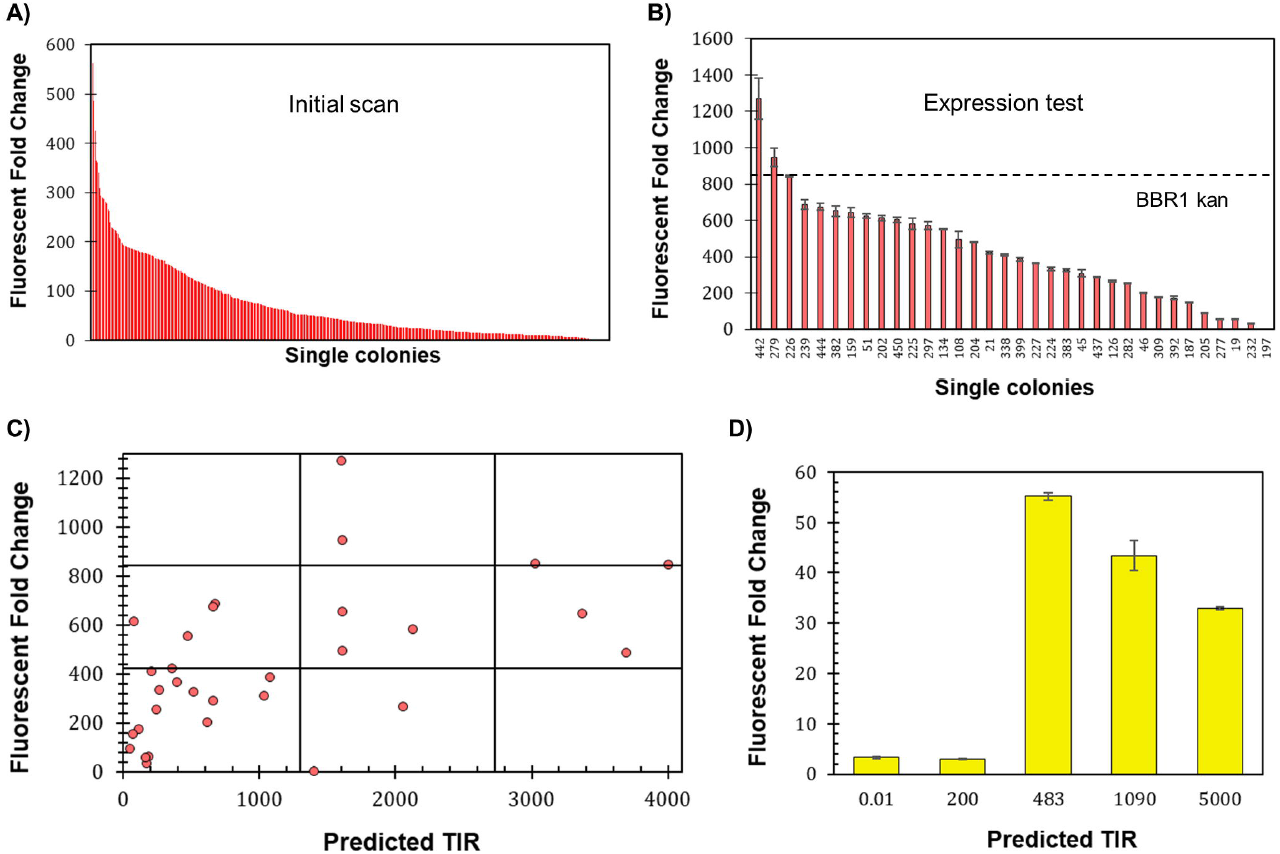
Determining 5’ UTR design guidelines for *R. palustris*. **A)** Fluorescent fold change scan of *R. palustris* BBR1 kan mRFP (RBS library) single colonies (Materials and Methods). **B)** Average fluorescent fold change of select *R. palustris* BBR1 kan mRFP (RBS library) strains grown in triplicate (Materials and Methods). Error bars indicate one standard deviation. **C)** Comparison of the predicted translation initiation rate (TIR) versus the fluorescent fold change average of the select *R. palustris* BBR1 kan (RBS library) mRFP strains. The grid lines divide each axis in thirds to aid in analysis. **D)** Average fluorescent fold change of *R. palustris* BBR1 kan (0.01, 200, 483, 1090, and 5000) 5’ UTR eYFP strains grown in triplicate (Materials and Methods). Error bars indicate one standard deviation.

**Figure 6.**
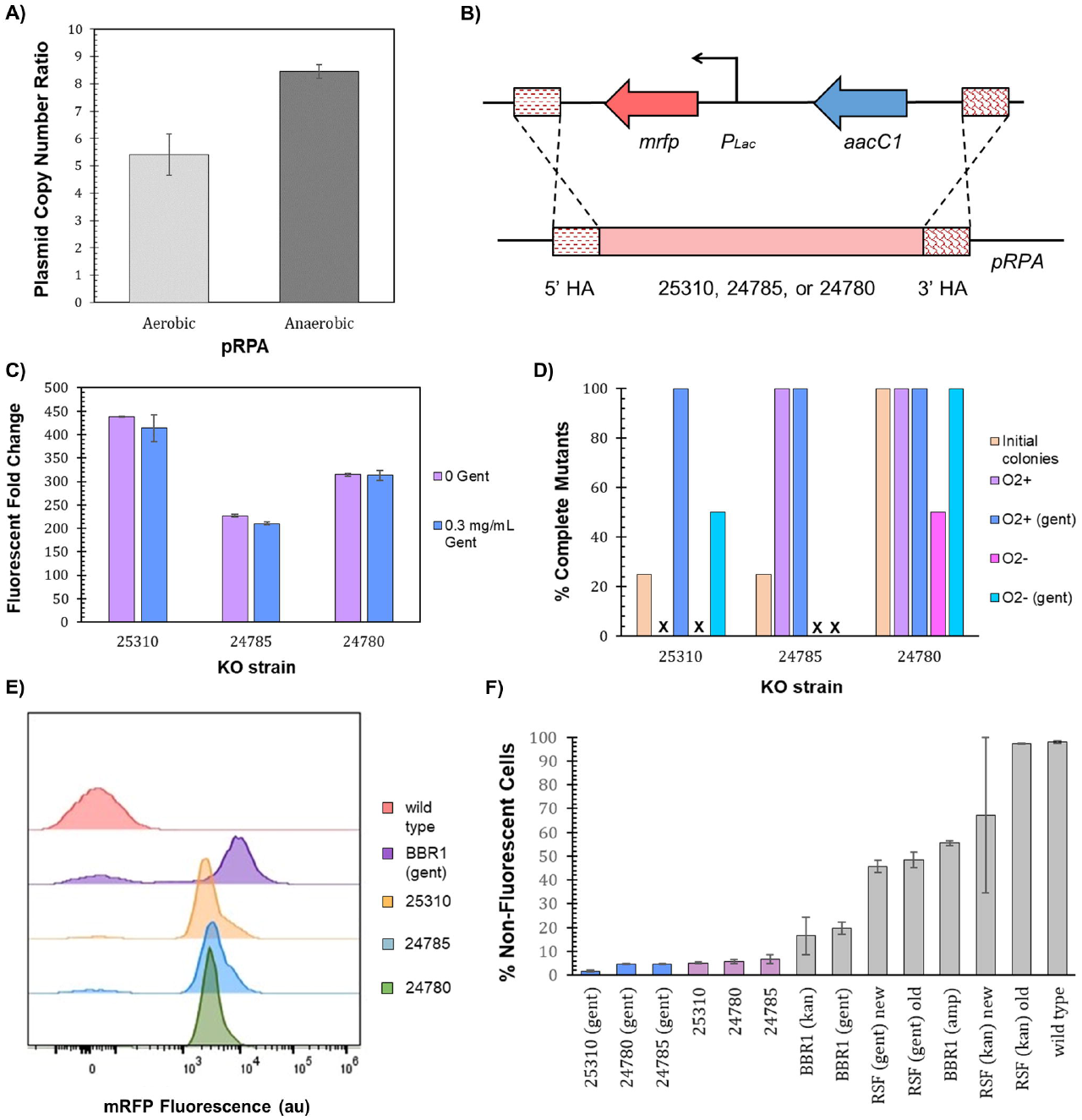
pRPA as a heterologous gene expression vector. **A)** Average pRPA copy number for wild type *R. palustris* grown aerobically in the dark and anaerobically in the light (100 μE white light) in duplicate. Copies of the plasmid’s replication protein gene (TX73_RS24750) was compared to the single copy *dxs* gene in the chromosome by qPCR and an external standard curve (Materials and Methods). Error bars indicate one standard deviation. **B)** Schematic of the *P_Lac_-mrfp* expression cassette and a gentamicin selection marker integrated into one of three locations, TX73_RS25310, TX73_RS24785, TX73_RS24780, in *R. palustris*’ endogenous plasmid, pRPA, by double homologous recombination. **C)** Fluorescent fold change average of *R. palustris* pRPA 25310 mRFP Gent, pRPA 24785 mRFP Gent, and pRPA 24780 Gent strains grown in triplicate either without any antibiotic or with 0.3 mg/mL gentamicin sulfate. The data is representative of the first test with new cultures as well as the test after three rounds of dilution and regrowth (Materials and Methods). Error bars indicate one standard deviation. **D)** Comparison of the percent complete mutants (those that do not produce an amplicon from PCR with primers binding to the DNA that is removed and also produce the correct sequence from PCR with primers outside of the homology arms) for the three pRPA locations. **E)** Representative flow cytometry results of mRFP fluorescence from *R. palustris* wild type, BBR1 gent mRFP, pRPA 25310 mRFP Gent, pRPA 24785 mRFP Gent, and pRPA 24780 Gent strains grown in triplicate. The growth procedure, media, and antibiotics are the same as for the fluorescent fold change test in Figure 6c (Materials and Methods). **F)** Average percent non-fluorescent cells for *R. palustris* strains grown in triplicate as determined by flow cytometry (Materials and Methods). Error bars indicate one standard deviation.

The optimize expression levels function of the DeNovo DNA RBS calculator was then used to design 5’ UTRs with higher (1090 and 5000) and lower (0.02 and 200) TIRs than the original pBBR1 kan eYFP plasmid, TIR of 483. The fluorescent fold change was then determined for the four subsequent strains following the same protocols used previously (Materials and Methods). The fluorescent fold change for the strains with high TIRs was lower than the original strain, 33-fold and 43-fold versus the original 55-fold (Figure 5d). The strains with the low TIRs produced very low fluorescent fold change, both approximately 3-fold.

### Harnessing pRPA for heterologous protein production

*R. palustris*’ active partitioning system ensures its endogenous plasmid, pRPA, is segregated into daughter cells (Debaugny et al., 2018), unlike exogenous plasmids (Meyer, 2009). Employing *R. palustris*’ endogenous plasmid could therefore address the problem of plasmid loss. pRPA encodes nine potential genes (Larimer et al., 2004) with five of the nine open reading frames annotated. Three of the open reading frames for hypothetical proteins are clustered together and one sits between genes for a replication protein and a protein associated with allocating the chromosomes and plasmids during cell division (Ebersbach and Gerdes, 2005). The copy number of pRPA was determined to average more than five copies per chromosome for aerobic cultures and more than eight for anaerobic cultures (100 μE white light) (Figure 6a) using qPCR and *R. palustris*’ gDNA as discussed earlier (Materials and Methods). In addition, there was minimal variation in the copy number for anaerobic cultures in contrast to the pBBR1 kan plasmid (Figure 3b).

Double homologous recombination with a p15A suicide plasmid (Materials and Methods) was then utilized to replace each of the three open reading frames that are clustered together (TX73_RS25310, TX73_RS24785, or TX73_RS24780) with a gentamicin selection marker and the *P_Lac_-mrfp* expression cassette, coded on the same DNA strand without a terminator between them (Figure 6b). After transformation with the suicide plasmids, four colonies were selected for each open reading frame replacement. The loss of the suicide plasmid and the incorporation of the expression cassette and selection marker was determined by colony PCR and sequencing of the PCR products. None of the colonies still harbored the suicide plasmid. One of four colonies for TX73_RS25310 and TX73_RS24785 (25310 and 24785) had completely integrated the expression cassette and selection marker (Figure 6d), as determined by the absence of a PCR product for primers that only bound to the DNA being replaced as well as an amplicon of the expected size and correct sequence for primers that bound outside of the homology arms. All four colonies for 24780 also showed complete segregation. The fluorescent fold change measurement protocol was then followed for cultures grown from a single colony that demonstrated complete segregation for each open reading frame replacement (Materials and Methods). The 25310 strain produced the highest fluorescent fold change, 400, while the fluorescent fold change for the 24785 and 24780 strains was 200 and 300 respectively (Figure 6c).

The stability of colonies with complete segregation was investigated by diluting the cultures just tested into media without gentamicin and also diluting the same culture into media with gentamicin, letting them reach stationary phase for at least 24 hours, and repeating the dilution and growth two more times. The presence or absence of gentamicin was maintained when diluting the last two rounds. Three rounds of dilution and growth were completed for cultures grown aerobically and one round was completed for cultures grown anaerobically (100 μE white light). Colony PCR, sequencing, and the fluorescence test was repeated at the end of the third aerobic round and the end of the first anaerobic round. There was no change in the fluorescent fold change, tested only in aerobic cultures (data not shown). The stability of the genetic change was dependent on the growth conditions, the presence of gentamicin in the media, and the open reading frame (Figure 6d). All strains maintained the expression cassette and gentamicin selection marker in some of copies of the plasmid; although, now a PCR product of the expected size for primers that only bound to the DNA being replaced was barely visible in the agarose gel for some of the cultures. Gentamicin ensured complete segregation for all cultures grown aerobically. Gentamicin maintained segregation for all of the anaerobically grown strain 24780 cultures and half of the strain 25310 cultures. Without gentamicin in the media, complete segregation was maintained in all of the strain 24780 and 24785 aerobic cultures and half of the strain 24780 anaerobic cultures.

Flow cytometry was then employed to further investigate the segregation of the expression cassette for all three strains, 25310, 24785, and 24780. Triplicates of each strain were grown with gentamicin in addition to triplicates of each strain without gentamicin, all in aerobic conditions for the flow cytometry test. Figure 6e is representative of the results. The averages and standard deviations are presented in Figure 6f. There was little variation in the number of cells with just the background fluorescence for the three strains utilizing pRPA, regardless of gentamicin in the media. The cultures of strain 24785 grown without gentamicin saw the largest average number of cells, 6.7%, with just the background fluorescence of the three strains and two growth conditions. The variation was also highest for strain 24785 grown without gentamicin, ±1.8%. This worst case pRPA mutant had a statistically lower number of cells with just the background fluorescence as compared to the most similar exogenous plasmid pBBR1 gent (student’s two-tail t-test, p < 0.05), which requires gentamicin.

## Discussion

*R. palustris*’ multiple modes of metabolism endow it with many valuable biochemical capabilities including utilizing energy from sunlight, fixing carbon, catabolizing recalcitrant aromatic compounds, and fixing nitrogen/producing hydrogen (Larimer et al., 2004). Furthermore, *R. palustris* can remain metabolically active in a non-growing state for months when supplied with just light and organic carbon (Gosse et al., 2010). Even with all of these beneficial traits, published work about engineering this microbe has been limited. *R. palustris*’ intrinsic resistance to antibiotics (Larimer et al., 2004) and the lack of basic synthetic biology tools that have been characterized for this PNSB are impediments to harnessing it’s potential. To address these problems, this work has focused on establishing a baseline understanding of the behavior of genetic parts (including selection markers, origins of replication, fluorescent reporters, terminators, and 5’ untranslated regions) that are fundamental to future engineering of this remarkable microorganism. Furthermore, this new knowledge provided the needed tools to take a significant step forward in establishing predictable heterologous gene expression in the chassis by employing its endogenous plasmid.

### Selection pressure

*R. palustris*’ antibiotic resistance makes the selection of a vector for gene expression in this bacterium critical. While the microbe was most sensitive to ampicillin, only 50% of the cells in the cultures were fluorescent at 50 μg/mL, the concentration that led to the best performance of the pBBR1 amp plasmid. The gene for a chloramphenicol acetyltransferase (CAT) is annotated in *R. palustris*’ genome and was expressed in both aerobic and anaerobic (100 μE white light) conditions. CAT attaches an acetyl group to the antibiotic, which prevents chloramphenicol from binding to the bacterium’s ribosome (Shaw, 1983). Removing *cat* did not increase *R. palustris*’ sensitivity to chloramphenicol, pointing to another mechanism that is responsible for the bacterium’s resistance. *R. palustris*’ intrinsic resistance to antibiotics has been attributed to its 22 unique resistance-nodulation-cell division (RND) pumps, which is more than has been discovered in any other bacterium (Larimer et al., 2004). RND pumps shuttle the substrate across the membrane and into the external medium, minimizing the internal substrate concentration. The energy source for RND pumps is a proton gradient (Fernando and Kumar, 2013), which could explain the increased instability of the pBBR1 and the pRPA knockout strains in anaerobic/photosynthetic conditions. These efflux pumps often have broad substrate specificity (Fernando and Kumar, 2013); in other words, one RND pump may be able to reduce the internal concentration of multiple antibiotics. Generally, these pumps are tightly regulated by interactions between local repressors and global regulators. As specific mechanisms of regulation are determined, possible solutions to the antibiotic resistance of a potential synthetic biology chassis could involve engineering the regulators to be less sensitive to the antibiotic(s) used for selection or altering the balance between the repressors and the inducers of the pumps.

Both kanamycin and gentamicin allowed heterologous protein production from pBBR1, but only gentamicin permitted stable maintenance of the RSF1010 plasmid. This might be related to gentamicin stopping growth of the wild type strain in 13 hours as opposed to more than 30 hours required for kanamycin. There were still 50% of the cells in the RSF1010 gent culture that only produced background fluorescence; although, that was stable over time. There is plenty of room for further investigation of selection markers, antibiotic concentrations, and plasmid stability in *R. palustris*. Spectinomycin allowed for even less growth than gentamicin, but over a significantly longer time. Spectinomycin’s resistance gene was therefore not tested for selection for this work, but it is a possibility in the future. Tetracycline was not even considered because of its light sensitivity, but maybe it could be used for selection in aerobic conditions. Higher concentrations of kanamycin and gentamicin could also be tested to see if plasmid stability improves. Although higher concentrations started to affect growth of *R. palustris* mutants, with a 17% decrease in growth at 600 μg/mL kanamycin or gentamicin as compared to wild type and a 26% decrease at 1200 μg/mL. Auxotrophies could also be investigated to identify a selection mechanism that does not involve antibiotics (Seif et al., 2020). If no auxotrophies are found in *R. palustris*, creating one through a knockout strain could be pursued.

### Origins of replication and plasmid stability

pBBR1 and RSF1010 are among the many origins of replication that have been identified as having a broad host range (Jain and Srivastava, 2013). While pBBR1 has been used for gene expression in *R. palustris* previously (Braatsch et al., 2006;Huang et al., 2010;Heiniger and Harwood, 2015), its stability and copy number had not been investigated in the microbe. RSF1010 belongs to a different incompatibility group than pBBR1 (del Solar et al., 1998;Meng et al., 2013). Its stability and copy number in *R. palustris* were also investigated since this would open the possibility of using the two plasmids together in the future. The origin of replication and selection marker both influenced plasmid stability, making it difficult to tease apart precise rules for use in this microbe. Both oris produced heterogeneous populations as shown by flow cytometry. It is not known from this data whether the non-fluorescent cells from all strains lost their plasmid or if the plasmid was present but mutated so that it no longer produced the fluorescent protein. A plasmid copy number of less than one per chromosome for the RSF1010 kan strain does suggest plasmid loss for at least some of the strains. Cell sorting could be used in the future to help answer this question. If the non-fluorescent cells for the RSF1010 gent strain lost their plasmid, our data suggests that it is not due to low plasmid copy number. The RSF1010 gent strain’s average plasmid copy was higher than that from the pBBR1 gent strain. In general, pBBR1sustained heterologous protein production with kanamycin selection, and both oris with gentamicin selection.

### Phototrophy and tool performance

To take advantage of the energy produced from *R. palustris*’ phototrophic (anaerobic) metabolism, the synthetic biology tools used to engineer the microbe also need to be characterized during this mode of growth. Due to the oxygen requirement of most fluorescent reporters and the bacterium’s background fluorescence, easy characterization through the use of a fluorescent reporter is not feasible. *mrfp* expression and plasmid copy number, both determined by qPCR, produced larger variation than what was seen from the same tests during aerobic growth. As discussed earlier, the proton gradient generated by photosynthesis might provide extra energy for RND pumps that are decreasing the selection pressure. Additionally, *R. palustris*’ photoheterotrophic metabolism is sensitive to how much light the microbe can absorb (Alsiyabi et al., 2019;Navid et al., 2019). Differences in light intensity are somewhat common throughout the growth chamber, which could lead to differences in the level of photosynthesis between the cultures in the chamber. Even with this variation, the pBBR1 kan strain results in anaerobic conditions were hopeful. The anaerobic culture with the lowest gene expression and plasmid copy number was well within the range of gene expression and copy number for the pBBR1 kan aerobic cultures.

### Transcriptional terminators

Terminators increase predictable gene expression by insulating the gene of interest from other nearby transcription units (Chen et al., 2013;Kelly et al., 2019), making them an important tool for regulating heterologous gene expression. Studies of terminator performance in *R. palustris* were not found. Furthermore, the construction of the pBBR1 kan mRFP (opposite strand) strain indicated that the original plasmid did not have an effective terminator between the selection marker and the expression cassette, *P_Lac_-mrfp*. Therefore, two terminators that had performed significantly different in *E. coli*, rrnC was 25 times stronger than tonB, were selected (Chen et al., 2013). Both terminators worked well on the pBBR1 kan mRFP plasmid in *R. palustris*, suggesting that other terminators characterized in *E. coli* would work well too. The difference in strength between the terminator, 75% versus 96%, was much less for *R. palustris* than for *E. coli*, serving as a reminder that genetic parts should be tested in the chassis of interest before designing expression vectors.

### Designing 5’ UTRs

Randomizing the six base pair ribosome binding site (RBS) for *mrfp* produced an expression cassette with a huge dynamic range. The question was whether or not the expression level could be predicted before the 5’ untranslated region (UTR) was changed. Choosing the expression level is important for balancing metabolic pathways and producing enzymes whose products are toxic at high concentrations. Comparing the translation initiation rates predicted by the RBS calculator to the normalized fluorescence for sequences across the dynamic range revealed that the level of predictability depended on the desired strength of expression. TIRs predicted to produce low expression generally produced low fluorescence. Once the predicted translation rate increased into the middle range, the fluorescent output was unpredictable. Sequences whose TIRs predicted a high level of expression all produced a lower level of fluorescence than was expected. The 5’ UTRs designed for *eyfp* fit into the same pattern. These revelations could aid in designing expression cassettes for *R. palustris* when the desired level of expression is known.

### Harnessing pRPA for heterologous protein production

Employing *R. palustris* endogenous plasmid for heterologous protein production could circumvent the plasmid loss that occurred with the pBBR1 kan strain. Furthermore, if complete segregation can be achieved selection pressure would not be needed, which is preferred for biotechnology applications. In addition, the plasmid copy number for pRPA showed little variation in aerobic and anaerobic conditions, which would improve the predictability of gene expression. pRPA’s plasmid copy numbers were also statistically similar to the plasmid copy numbers for the pBBR1 plasmids from cultures grown aerobically (student’s two-tail t-test, p > 0.05). This indicates that pRPA should be a good alternative to exogenous plasmids for heterologous gene expression.

During aerobic growth, more than 90% of the cells in the pRPA cultures for all three open reading frames were fluorescent, as compared to 80% for the BBR1 gent strain. The level of normalized fluorescence was dependent on the open reading frame that was replaced, but not the presence of gentamicin in the media, even across the span of three rounds of dilution and regrowth. The open reading frame also affected segregation. The expression cassette was maintained in at least some of the copies of pRPA for all constructs in all conditions (oxygen and gentamicin), but there were instances of the original DNA sequence being detected by PCR for at least one condition for all three open reading frames at the completion of testing. This occurred after all three strain’s original colony demonstrated complete segregation. At the completion of testing, the 25310 strain required gentamicin to demonstrate complete segregation in all cultures. Strain 24785 only demonstrated complete segregation when grown aerobically. Strain 24780 was most successful, with complete segregation present in all but half of the cultures grown anaerobically without gent. Interestingly, the *R. palustris* Δ*cat* strain constructed through sucrose counterselection also produced PCR products, or the lack thereof, that indicated complete segregation, but when a double knockout was attempted the original *cat* sequence that was removed was now detectable by PCR. No reports of this behavior were found in literature. Whether ploidy could be playing a role is under investigation.

Stable and predictable heterologous protein production can be achieved in the metabolically robust *R. palustris* when the genetic parts have been well characterized for the condition(s) of interest and their limitations are understood. It was not intuitive for the pink bacterium’s background fluorescence to be lowest for mRFP’s excitation and emission wavelengths. The 5’ untranslated regions functioned within a reasonable range, as long as the predicted translation initiation rate was for a low level of expression. The two transcriptional terminators did a good job of insulating the expression cassette; although, their level of performance could not be predicted from studies in *E. coli*. Both of the pBBR1 and RSF1010 plasmids were stable over time with gentamicin selection, but only the pBBR1 plasmid was stable with the kanamycin selection marker. Furthermore, gene expression, exogenous plasmid copy number, and the stability of genetic changes were more consistent for aerobic conditions than for anaerobic conditions; yet, there is a potential for high levels of protein production when the bacterium is performing photosynthesis. Finally, utilizing pRPA as an expression vector shows promise. Significant levels of mRFP were generated from the endogenous plasmid, and two of the integration sites did not require selection pressure in aerobic conditions after strain validation. While not all of *R. palustris*’ secrets have been unlocked, this work has begun to clarify rules for expressing heterologous genes in the complex microbe, laying the groundwork for future engineering endeavors that harness its unique biochemical potential.

## Acknowledgements

We gratefully acknowledge funding support from National Science Foundation (NSF) CAREER grant (25-1106-0039-001) awarded to R.S. and from the U.S. Department of Agriculture, National Institute of Food and Agriculture (USDA-NIFA) Postdoctoral Fellowship 2019-67012-29632 awarded to C.I. We would like to thank Dr. Himadri Pakrasi and The Pakrasi Lab (Washington University in St. Louis) for the donation of pSL3068. We would also like to thank the Flow Cytometry Core Facility in the University of Nebraska – Lincoln’s Center for Biotechnology for the flow cytometry measurements, protocol development, data analysis, and graph creation.

## Author Statement

CI and MK contributed equally to this work. CI and RS formulated the idea for this work. CI and TC developed the methodology. CI, MK, and TC constructed strains and conducted tests. All authors analyzed the data. CI and MK wrote the manuscript with guidance from RS.

## Supplemental Material

Supplementary Tables I-VI and Supplementary Figures 1-4 as described in the text.

