## Supplemental Material for "Synthetic Biology Tool Development Advances Predictable Gene Expression in the Metabolically Versatile Soil Bacterium *Rhodopseudomonas palustris*"

**Supplementary Table I.** Plasmids/Strains used in this work.

| **Name** | **Parts** | **Type and Source** |
| --- | --- | --- |
| pJQ200SK | p15A ori; *gent^R^; P_sacb_:sacb* | Integrative  (Quant, Hynes, 1993) |
| pBBR1MCS-2 | pBBR1 replicon; *kan^R^*; *P_Lac_* | Replicative  (Kovach et al., 1995) |
| pBbB7a-GFP | *P_amp:_amp^R^; gfpuv* | Replicative  (Lee et al., 2011) |
| pSL3068 | RSF1010 replicon; *eyfp* | Replicative  (Knoot et al., 2019) |
| pCMrfp | *mrfp* | Replicative  (Bi et al., 2013) |
| pRPA | *R. palustris* endogenous plasmid | Replicative |
| pΔcat | p15A ori; *5’HA CAT; 3’HA CAT; gent^R^; P_sacb_:sacb* | Integrative  This Study |
| BBR1 kan GFPuv | pBBR1 replicon; *kan^R^; P_Lac_:gfpuv* | Replicative  This Study |
| BBR1 kan 483 eYFP | pBBR1 replicon; *kan^R^; P_Lac_:eyfp* | Replicative  This Study |
| BBR1 kan mRFP | pBBR1 replicon; *kan^R^; P_Lac_:mrfp* | Replicative  This Study |
| BBR1 kan mRFP (opposite strand) | pBBR1 replicon; *kan^R^; P_Lac_:mrfp* | Replicative  This Study |
| BBR1 amp mRFP | pBBR1 replicon; *P_amp_:amp^R^; P_Lac_:mrfp* | Replicative  This Study |
| BBR1 gent mRFP | pBBR1 replicon; *gent^R^; P_Lac_:mrfp* | Replicative  This Study |
| BBR1 kan mRFP tonb | pBBR1 replicon; *kan^R^; ton B; P_Lac_:mrfp* | Replicative  This Study |
| BBR1 kan mRFP rrnc | pBBR1 replicon; *kan^R^; rrnc; P_Lac_:mrfp* | Replicative  This Study |
| BBR1 kan 0.01 eYFP | pBBR1 replicon; *kan^R^; P_Lac_:0.01 5’ UTR eyfp* | Replicative  This Study |
| BBR1 kan 200 eYFP | pBBR replicon; *kan^R^; P_Lac_:200 5’ UTR eyfp* | Replicative  This Study |
| BBR1 kan 1090 eYFP | pBBR replicon; *kan^R^; P_Lac_:1090 5’ UTR eyfp* | Replicative  This Study |
| BBR1 kan 5000 eYFP | pBBR1 replicon; *kan^R^; P_Lac_:5000 5’ UTR eyfp* | Replicative  This Study |
| pRPA 24780 mRFP Gent | P15A ori; *3’HA 24780; gent^R^; P_Lac_:mrfp; 5’HA 24780* | Integrative  This Study |
| pRPA 24785 mRFP Gent | p15A ori; *3’HA 24785; gent^R^; P_Lac_:mrfp; 5’HA 24785* | Integrative  This Study |
| pRPA 25310 mRFP Gent | p15A ori; *3’HA 25310; gent^R^; P_Lac_:mrfp; 5’HA 25310* | Integrative  This Study |

**Supplementary Table II.**  List of genetic parts used in this work.

| Part name | Type and source | DNA sequence |
| --- | --- | --- |
| *amp^R^* | Antibiotic resistance gene  (Lee et al., 2011) | atgagtattcaacatttccgtgtcgcccttattcccttttttgcggcattttgccttcctgtttttgctcacccagaaacgctggtgaaagtaaaagatgctgaagatcagttgggtgcacgagtgggttacatcgaactggatctcaacagcggtaagatccttgagagttttcgccccgaagaacgttttccaatgatgagcacttttaaagttctgctatgtggcgcggtattatcccgtattgacgccgggcaagagcaactcggtcgccgcatacactattctcagaatgacttggttgagtactcaccagtcacagaaaagcatcttacggatggcatgacagtaagagaattatgcagtgctgccataaccatgagtgataacactgcggccaacttacttctgacaacgatcggaggaccgaaggagctaaccgcttttttgcacaacatgggggatcatgtaactcgccttgatcgttgggaaccggagctgaatgaagccataccaaacgacgagcgtgacaccacgatgcctgtagcaatggcaacaacgttgcgcaaactattaactggcgaactacttactctagcttcccggcaacaattaatagactggatggaggcggataaagttgcaggaccacttctgcgctcggcccttccggctggctggtttattgctgataaatctggagccggtgagcgtgggtctcgcggtatcattgcagcactggggccagatggtaagccctcccgtatcgtagttatctacacgacggggagtcaggcaactatggatgaacgaaatagacagatcgctgagataggtgcctcactgattaagcattggtaa |
| *kan^R^* | Antibiotic resistance gene  (Kovach et al., 1995) | atgattgaacaagatggattgcacgcaggttctccggccgcttgggtggagaggctattcggctatgactgggcacaacagacaatcggctgctctgatgccgccgtgttccggctgtcagcgcaggggcgcccggttctttttgtcaagaccgacctgtccggtgccctgaatgaactgcaggacgaggcagcgcggctatcgtggctggccacgacgggcgttccttgcgcagctgtgctcgacgttgtcactgaagcgggaagggactggctgctattgggcgaagtgccggggcaggatctcctgtcatctcaccttgctcctgccgagaaagtatccatcatggctgatgcaatgcggcggctgcatacgcttgatccggctacctgcccattcgaccaccaagcgaaacatcgcatcgagcgagcacgtactcggatggaagccggtcttgtcgatcaggatgatctggacgaagagcatcaggggctcgcgccagccgaactgttcgccaggctcaaggcgcgcatgcccgacggcgaggatctcgtcgtgacccatggcgatgcctgcttgccgaatatcatggtggaaaatggccgcttttctggattcatcgactgtggccggctgggtgtggcggaccgctatcaggacatagcgttggctacccgtgatattgctgaagagcttggcggcgaatgggctgaccgcttcctcgtgctttacggtatcgccgctcccgattcgcagcgcatcgccttctatcgccttcttgacgagttcttctga |
| *gent^R^* | Antibiotic resistance gene  (Quandt and Hynes, 1993) | atgttacgcagcagcaacgatgttacgcagcagggcagtcgccctaaaacaaagttaggtggctcaagtatgggcatcattcgcacatgtaggctcggccctgaccaagtcaaatccatgcgggctgctcttgatcttttcggtCgtgagttcggagacgtagccacctactcccaacatcagccggactccgattacctcgggaacttgctccgtagtaagacattcatcgcgcttgctgccttcgaccaagaagcggttgttggcgctctcgcggcttacgttctgcccaagtttgagcagccgcgtagtgagatctatatctatgatctcgcagtctccggcgagcaccggaggcagggcattgccaccgcgctcatcaatctcctcaagcatgaggccaacgcgcttggtgcttatgtgatctacgtgcaagcagattacggtgacgatcccgcagtggctctctatacaaagttgggcatacgggaagaagtgatgcactttgatatcgacccaagtaccgccacctaa |
| *tonB* | Terminator  (Postle and Good, 1983) | agtcaaaagcctccggtcggaggcttttgact |
| *rrnC* | Terminator  (Young, 1979) | cagataaaaaaaatccttagctttcgctaaggatgatttct |
| *P_Lac_* | Promoter  (Kovach et al., 1995) | ggcagtgagcgcaacgcaattaatgtgagttagctcactcattaggcaccccaggctttacactttatgcttccggctcgtatgttgtgtggaattgtgagcggataacaat |
| *P_SacB_* | Promoter  (Quant, Hynes, 1993) | cacatatacctgccgttcactattatttagtgaaatgagatattatgatattttctgaattgtgattaaaaaggcaactttatgcccatgcaacagaaactataaaaaatacagagaatgaaaagaaacagatagattttttagttctttaggcccgtagtctgcaaatccttttatgattttctatcaaacaaaagaggaaaatagaccagttgcaatccaaacgagagtctaatagaatgaggtcgaaaagtaaatcgcgcgggtttgttactgataaagcaggcaagacctaaaatgtgtaaagggcaaagtgtatactttggcgtcaccccttacatattttaggtctttttttattgtgcgtaactaacttgccatcttcaaacaggagggctggaagaagcagaccgctaacacagtacataaaaaaggagacatgaacg |
| *P_amp_* | Promoter  (Lee et al., 2011) | cgcggaacccctatttgtttatttttctaaatacattcaaatatgtatccgctcatgagacaataaccctgataaatgcttcaataatattgaaaaaggaagagt |
| *sacb* | Levansucrose gene  (Quant, Hynes, 1993) | atgaacatcaaaaagtttgcaaaacaagcaacagtattaacctttactaccgcactgctggcaggaggcgcaactcaagcgtttgcgaaagaaacgaaccaaaagccatataaggaaacatacggcatttcccatattacacgccatgatatgctgcaaatccctgaacagcaaaaaaatgaaaaatatcaagttcctgaattcgattcgtccacaattaaaaatatctcttctgcaaaaggcctggacgtttgggacagctggccattacaaaacgctgacggcactgtcgcaaactatcgcggctaccacatcgtctttgcattagccggagatcctaaaaatgcggatgacacatcgatttacatgttctatcaaaaagtcggcgaaacttctattgacagctggaaaaacgctggccgcgtctttaaagacagcgacaaattcgatgcaaatgattctatcctaaaagaccaaacacaagaatggtcaggttcagccacatttacatctgacggaaaaatccgtttattctacactgatttctccggtaaacattacggcaaacaaacactgacaactgcacaagttaacgtatcagcatcagacagctctttgaacatcaacggtgtagaggattataaatcaatctttgacggtgacggaaaaacgtatcaaaatgtacagcagttcatcgatgaaggcaactacagctcaggcgacaaccatacgctgagagatcctcactacgtagaagataaaggccacaaatacttagtatttgaagcaaacactggaactgaagatggctaccaaggcgaagaatctttatttaacaaagcatactatggcaaaagcacatcattcttccgtcaagaaagtcaaaaacttctgcaaagcgataaaaaacgcacggctgagttagcaaacggcgctctcggtatgattgagctaaacgatgattacacactgaaaaaagtgatgaaaccgctgattgcatctaacacagtaacagatgaaattgaacgcgcgaacgtctttaaaatgaacggcaaatggtacctgttcactgactcccgcggatcaaaaatgacgattgacggcatttcgtctaacgatatttacatgcttggttatgtttctaattctttaactggcccatacaagccgctgaacaaaactggccttgtgttaaaaatggatcttgatcctaacgatgtaacctttacttactcacacttcgctgtacctcaagcgaaaggaaacaatgtcgtggtgattacaagctatatgacaaacagaggattctacgcagacaaacaatcaacgtttgcgccaagcttcctgctgaacatcaaaggcaagaaaacatctgttgtcaaagacagcatccttgaacaaggacaattaacagttaacaaataa |
| *eyfp* | Fluorescent reporter gene  (Knoot et al., 2019) | atggtgagcaagggcgaggagctgttcaccggggtggtgcccatcctggtcgagctggacggcgacgtaaacggccacaagttcagcgtgtccggcgagggcgagggcgatgccacctacggcaagctgaccctgaagttcatctgcaccaccggcaagctgcccgtgccctggcccaccctcgtgaccaccttcggctacggcctgcaatgcttcgcccgctaccccgaccacatgaagctgcacgacttcttcaagtccgccatgcccgaaggctacgtccaggagcgcaccatcttcttcaaggacgacggcaactacaagacccgcgccgaggtgaagttcgagggcgacaccctggtgaaccgcatcgagctgaagggcatcgacttcaaggaggacggcaacatcctggggcacaagctggagtacaactacaacagccacaacgtctatatcatggccgacaagcagaagaacggcatcaaggtgaacttcaagatccgccacaacatcgaggacggcagcgtgcagctcgccgaccactaccagcagaacacccccatcggcgacggccccgtgctgctgcccgacaaccactacctgagctaccagtccgccctgagcaaagaccccaacgagaagcgcgatcacatggtcctgctggagttcgtgaccgccgccgggatcactctcggcatggacgagctgtacaaggctgcgaattga |
|  | 5’ 0.01 UTR  5’ 200 UTR  5’ 483 UTR (Original)  5’ 1090 UTR  5’ 5000 UTR | ttaggtcgaattcaggtaccatcacctgcatcgttac  gttgttaagtcaataaaggtaccatttggt  tttcaggaattcaaaagatcttttaagaaggagatatacat  ttttcgatcaaacatagtaggaggtctcct  ttaattttgtctggttagattagggggtttttag |
| *mrfp* | Fluorescent reporter gene  (Bi et al., 2013) | atgagtaaaggagaagaacttttcactggagttgtcccaattcttgttgaattagatggtgatgttaatgggcacaaattttctgtcagtggagagggtgaaggtgatgcaacatacggaaaacttacccttaaatttatttgcactactggaaaactacctgttccgtggccaacacttgtcactactttctcttatggtgttcaatgcttttcccgttatccggatcacatgaaacggcatgactttttcaagagtgccatgcccgaaggttatgtacaggaacgcactatatctttcaaagatgacgggaactacaagacgcgtgctgaagtcaagtttgaaggtgatacccttgttaatcgtatcgagttaaaaggtattgattttaaagaagatggaaacattctcggacacaaactggagtacaactataactcacacaatgtatacatcacggcagacaaacaaaagaatggaatcaaagctaacttcaaaattcgccacaacattgaagatggctccgttcaactagcagaccattatcaacaaaatactccaattggcgatggccctgtccttttaccagacaaccattacctgtccacacaatctgccctttcgaaagatcccaacgaaaagcgtgaccacatggtccttcttgagtttgtaactgctgctgggattacacatggcatggatgagctctacaaataa |
| *gfpuv* | Fluorescent reporter gene  (Lee et al., 2011) | atgagtaaaggagaagaacttttcactggagttgtcccaattcttgttgaattagatggtgatgttaatgggcacaaattttctgtcagtggagagggtgaaggtgatgcaacatacggaaaacttacccttaaatttatttgcactactggaaaactacctgttccgtggccaacacttgtcactactttctcttatggtgttcaatgcttttcccgttatccggatcacatgaaacggcatgactttttcaagagtgccatgcccgaaggttatgtacaggaacgcactatatctttcaaagatgacgggaactacaagacgcgtgctgaagtcaagtttgaaggtgatacccttgttaatcgtatcgagttaaaaggtattgattttaaagaagatggaaacattctcggacacaaactggagtacaactataactcacacaatgtatacatcacggcagacaaacaaaagaatggaatcaaagctaacttcaaaattcgccacaacattgaagatggctccgttcaactagcagaccattatcaacaaaatactccaattggcgatggccctgtccttttaccagacaaccattacctgtccacacaatctgccctttcgaaagatcccaacgaaaagcgtgaccacatggtccttcttgagtttgtaactgctgctgggattacacatggcatggatgagctctacaaataa |

**Supplementary Table III.** Flanking regions used for sucrose counter-selection and double homologous recombination

| **Name** | **DNA sequence** | **Location** |
| --- | --- | --- |
| 5’ HA CAT | gagtccttcttcacatacagcaccgagtgatagcccttggagccgtcaccgttgacctcgatggcgaacggctcgaccttggcgccgaccagataggcgcgggcgtaggacgccgggccgtacatgccgatgtggatgttgccggcgcgctggccttcgatcaccgcggcgtagtcgttggcgatgcggagtttcaccggcacgccgagctgcttggtgaggtagtcggtcagcggggtccagcgggtggtggtgcccgaggcgttctcgtccgggatcttgccgaacaccagttcgggatatttcgccttccagtcctgcgccggcgcggcgtgcgccgtgaacgccagcgcggcagcggcggcgacgagagtacggagcttgatcatgacgtttcctgtctcttcaagtcgatgaacggtcgaaacacggcaaccgcacgcggtcgcgccgcacgcggtgccggataggggagacggcgaacgatcgccgccgggatcaggccacggcgacgccggccaatgcgggggcgggaaggccgtgcggcacgtgcgacgcggtgccgagtacttcgtcggcttcgaggtcgtagagttcgcgcgcgatctggtcggtgagcgcggccggcgcgccatcgaacaccacgcggccggaggccatgccgatcagtcggtcgcaatagctgcgggccagatcgagcgagtgcagattgcacagcacggtgatgccgaagtgcttgttgatgcgcagcagcgcatccatcacgatcttggtgttgcgcggatcgagcgaggcgatcggctcgtcggcgagcacgatatcgggctgctgcaccagggcgcgggcgatcgcgacgcgctgctgctgaccgccggagagctggtcggcgcgctgcgccgcgtaggacgccatgtcgaactgatcgagcgccgagatcgccagggcgcgatcctgctcgggccacatctgcaccagcgagcgccacgacggcacttcggacagccggcccatcagcacgttggtcagcacgtcgagccggccgatcaggttgaattgctggaagatcatcgccgagcgtgcgcgccattgccgcagctcgcggccctgcagcgcggtcacatcaataccttcgaacaggatgcggccctcggaaggctccgccagccggttcagcatgcgcagcagagtcgacttgcctgcgccggatcggccgatcaccccgacgaagctgccgcgctccaccgaaaacgatgcgccatctacggcggctttgctgccaaaacggcaggtcagaccctcaactaccagcatgcgctgctcccgaaacaggttcgggaggaatcgctaacgccgttgttttgcaaccgtgtgacagtcgatgtagttccgccgacatatcaaggtcttcgctacgtcatcgatgcgtcacgacgacgtcatgagtggcaccgatgacaggctcaactccaatcatcgcgcacgagaatcccatg | chromosome |
| 3’ HA CAT | aggaaactgtcagtggacgaattcctcgacagatacgaagctgcggccaaggcgcagccggcggacgcccgtgtggctgtaacggacgagcaggtgatgggatgagcgagatcgcgatcgagggcggccgcacgctggtcggcgacgagattgccgagggctcgctgcagatcgctggcggcgcgatcgcggtggtcggttcggcgagcggtcaggcggcgatccggatcgacgcccgcggcctcttggtgttgcccggcatcgtcgatctgcacggcgatgcgttcgagcggcagatgatgccgcggccgggcgtcgatttcccgatcgacgtggcgctgatcgacagcgaccgccaggcgatcgctaacggtctgacgacggtctttcatgccgtcacctggtcgtgggagcccggcctgcgcagtgccgacaacgctcgccggatgctcgacgcgatcgatctgaccaagccgcggctttccgccgacacccgcattcatctgcgccacgaaacctttaatctcgatgcggtcgacgagatcgcgcaatggatcggcgaccgccgcgtcgatctgttcgccttcaacgatcacatggacacgacgctggcgaacctcgccaagccgcagaagcggaaccggatggtcgagcgcaccggtctgtccgacgacgaattcgaccggctggtcgaacgcatcgcggcgcgtggcgacgaagtgccacaggcaagtcggcgcctcgccgccatcgcacgagaggtgtcgatgccgatgctgtcgcacgatgacgacacgccggcgatacggcagggctaccgtgatctcggcgtcggcatcgccgaattccccaccaccgaggagaccgcacgcgacgcggcgctgcacggcgacttcatcgtgttcggctcgccgaacgtggtgcgcggtggcagccacaccggctggaccaaggcggccgacatgatcgccaagggactgtgctcggtgctggcctccgactactactacccggcgcccctgctcgccgcgttccggctggtgcatgacggcatcctgccgctggagcaggcctggaagttgatctcggagaaccccgcccgcgccgctggtctgtctgaacgcggcacgatcgcgagcggtcagcgcgccgacgtgctgctggtcgacgatcgcgtcgcgctgcggccgcggatcgtcgcggtgatcgccggcggccggctggtgcacgtcgacgatccggagcggttcgtgatcacgcgcaaactgccgcacccggctggcgttgccgcttgatctgacgttatcatcatcgg | chromosome |
| 5’ HA 27480 | ttgttgcggtcattggctcaaattcgatctcaggcccgaatcgtaacgcagcctgatgcttttggcaccaaacccacaaagaatcctgtgcatctgcgtggctatttcgtcgaaggggcagaaaccccgttcgggagcctgctgaatcggcggttaggaggcggcgttcgtctttagcctggaaatcgtcggcgggctgacgttgaaaatcggggcgatctcgcggacgcctcgccctgcggccagcatcgcgcgggcttcgtcttgctgggctggggtgagcttgtagggtcggcccaggctcttgcccttcgccttggcgcgggcgcggccttctgacgtgcgggcgaggatggtggcccgttccagcttggcggcggcgcctagcacggtcagcatgaactcggccagctcggtgctggtatcggcccaaggctcggcgatggatttgaaggttgccccggcctccttcacatcgtgcacgatgttcagcaagtcgcgggtggagcgcgcgaggcggtcgatgcgcgtgaccagcagcacgtcccccgcgtccagggcggcgagggccttcttgagctgggggcgctcggcgctggcgccgctaatgttctcgcgaaacaccttggccgctcctgcggccgtgagctgctcgacttgggcggccgtgtcctggtcactggtggagacgcgcgcgtagccgtaaatcatgccgcattgtgtaactgacttatgaaacggtcaagaggcttgatgcagttggatttttctttgtttcataagttttgacttatgaactaacccggctggggaaagcgggtatggcggtttgttgcccttccgttcatttcacgactttcgtataagattcgggttcctccccacttcatttagtaatagtggccgcgacatgcccaaaatcttgc | Endogenous plasmid  pRPA |
| 3’ HA 27480 | gtttcgcctcatcttcccggaatctaaagccgcgttgcacccttcgcttccgttccctccaatttgcaccgaaccggcggcataaatgctggcggtgctaatcatgcctgtggtcgagatgcccggtccgaccaggacgcccgtcgttccgaaataggtggtgccactcaccccgcctgtggtgaagctgatggtgccaccattccccgccaccacatttgcggaggtgctgacgatgcggtcgccgctgttgctgccggtgatggcgacgccattgacggtgagcgcggtcgcgctgatgttggtggccgtgaggcggctgctggcgttggtccaggtgaggccactgttggcatcgaagctgttgcccgtgttgtactggatatagccggtgctgcccgcggccacggtgctgccgccgcttagcgggttgttccagccgttgctggtgccgttgaaaacgcagaaggtgacggcaacgttctccggccgcaagcatattggcgagcgcctggctctcctggtgagggccaactacgtcgctcgggcacgcagcttggcggcccgccccggcgtgcactggctgacgccaaaggcagccacgcacttccaccctgtctttggtgaggaatgggcgcgcgcggcgagtggcaaggggtttacggataacctgcatatgccgcagcggctgctggactacgccatgggccgccagaagggcagcgccgcgacccgcatgatgttcgtgtttggcagcgcggcaatgttagagcgcgccatgaccaccctgcccgaccgcgacgctcgggtctggcgcgcggtgttcttcaaatccttgccggacttggtagaagattttggcgggacctggttgcaggcaactggcgagcgccggaatccgttcgagcgggtggaacctgcccaagcgacaggagccccatagagacgtttttgggggaggtgccgagcgttcccttgctcgcggcgttttaatcgctgtggtggccgctggggctgttctgtgtcgccttggatcaaaggcgaggtccgcctttgtccatggcagccacttt | Endogenous plasmid  pRPA |
| 5’ HA 24785 | ccgcattgtgtaactgacttatgaaacggtcaagaggcttgatgcagttggatttttctttgtttcataagttttgacttatgaactaacccggctggggaaagcgggtatggcggtttgttgcccttccgttcatttcacgactttcgtataagattcgggttcctccccacttcatttagtaatagtggccgcgacatgcccaaaatcttgcgggaaaccgcacatatcagccggaagctgacgcagaacatctttcgagccttcgactatgcgcgtgcaggtgggtttccactcaacctttacgttgtcatcaacattcgggaaactgacgcctgcgccgcagcttcggcctttgaacgtatccgccacaaatatcgggactggttggcccaccactctcgcaagctcggggtccgcatcccgcctatgtacgtgttcacctttgaggcgccaggccatccgcatgtgaactgggcgttgcgcgtcccgccgcgcctggttgatgagtttcagcggaaactgcccggctgggtagagaaggtgcaggggccactaggtccctttgacatcaacgtgcagccgatagcgccggacggcgcatataaggcgctggccaactacatcgtaaaaggctgtgacccggagtatgtggcgcactttcatctcgcagcccttgccgaacagcatggcccgcagggtgcgttttggggaaggcgggcaggggttagcccgtcgctcaacaaagcagagcgagacgcggccggattcaatccaaaacgccgggaggtgcgcagccgctctcatgaacgggacgccgcgtgaatgctacctgcaaacctgcatatgcccggtcgtggggtttcgcctcatcttcccggaatctaaagccgcgttgcacccttcgcttccgttccctccaatttgcaccgaaccggcggcataaatgctggcggtgctaatcatg | Endogenous plasmid  pRPA |
| 3’ HA 24785 | gacaggagtttagccatgggattagtcagagtccctagccattgctgatgcgagattatttaaactttgccaagggatattgaccagacgggccgtgctaccaagccgttgctgtcgcaagtgtcgcttaacccgcttctgggctatccaatggttattaatcttgtaaacgcttatcctgccctgctcactggcaatgtagaaccgcttatggactctaagccagttcttccagtttggactctccaaactcagagggtacttagtccctgcaccgtgcaagtatccgtcaatcaccgtcggttgaggctccatacctacattgcccctatcaagccagttttagcacgctgggaagcacgataatctactgcgactaccctagggtactcccgtagaaaagcgttatgaagtatgaccggcaaaaaaccacattgaccctatcagagcaaagctagcacggtagggaacgcgataatctactgcaacacctataggtctgttgctgacacgcacgatgacgtatgacccttttagcacggtagggagcgtgataatcttctgcaacacctatatagtattgttgcgatcgcgagcgatggcgt | Endogenous plasmid  pRPA |
| 5’ HA 25310 | gcctggttgatgagtttcagcggaaactgcccggctgggtagagaaggtgcaggggccactaggtccctttgacatcaacgtgcagccgatagcgccggacggcgcatataaggcgctggccaactacatcgtaaaaggctgtgacccggagtatgtggcgcactttcatctcgcagcccttgccgaacagcatggcccgcagggtgcgttttggggaaggcgggcaggggttagcccgtcgctcaacaaagcagagcgagacgcggccggattcaatccaaaacgccgggaggtgcgcagccgctctcatgaacgggacgccgcgtgaatgctacctgcaaacctgcatatgcccggtcgtggggtttcgcctcatcttcccggaatctaaagccgcgttgcacccttcgcttccgttccctccaatttgcaccgaaccggcggcataaatgctggcggtgctaatcatgcctgtggtcgagatgcccggtccgaccaggacgcccgtcgttccgaaataggtggtgccactcaccccgcctgtggtgaagctgatggtgccaccattccccgccaccacatttgcggaggtgctgacgatgcggtcgccgctgttgctgccggtgatggcgacgccattgacggtgagcgcggtcgcgctgatgttggtggccgtgaggcggctgctggcgttggtccaggtgaggccactgttggcatcgaagctgttgcccgtgttgtactggatatagccggtgctgcccgcggccacggtgctgccgccgcttagcgggttgttccagccgttgctggtgccgttgaaaacgcagaaggtgacggcaacgttctccggccgcaagcatattggcgagcgcctggctctcctggtgagggccaactacgtcgctcgggcacgcagcttggcggcccgccccggcgtgcactggctgacgccaaaggcagccacgcacttccaccctgtctttggtgaggaatgggcgcgcgcggcgagtggcaaggggtttacggataacctgcatatgc | Endogenous plasmid  pRPA |
| 3’ HA 25310 | ctttgtccatggcagccactttgcactgcctgagctgcagtcaagggcgcgaagcgccgaagggaacccttgacggcttaggcgagggctgtgcctccccctttttcttcaaatccaattttcagaatctaccgccgaaggcgttttaacaagtttctatagttttaagtaagttttaggggtgaaatttcccttttgaatcaaggtggttagagggtggttttgtccgcgatagacggtctgatgtccgtgatagacggttttttgtccgcgatagacggtctcctgtccgcgatagacgttcgggccaaaagttatccacaggccggaatctctgggtttaagtggctgaaaacatgttgttgcgggcgatagacggcttggccggaattgtccgcgatagacggtcgggccaaaaatccgggccaaaattaattggccgcgagctgaatgacttccggcgtggtcggcatcgcgggccgccgcttcataagcaaaatcggccctcccttcccttgctctagcgccaggtcaaaatctggcatcgggaaagtctgcgtgcgggtcacaaactccttcaggtcgaaaaggaacttgcgcggctcccgtgacgagccgctgcgcttgtgaagttcggcgacggaatatcgcgcctcgtccttccccgccgccttgcgggccagccgatagataaaccggccgaggcccgacgtgatgaggaagtaatccgggtgcagagtgagcaagggcaacgccttgtcggatctcacgacgctcgaataaacccagtcggggatgcggatttcgatctggtccaccttgccggtgttggtggagctgatgatggtgtgttcaccaataaggggccgggtatcaacctggcggcgctttccgcccgcaaggtttgtgatggtgatggaggttttcgataaccggcgcagggcggcttcaatctcttgatattgtcgtccgccgctggcgcggcggctgaacttcaaaagatgcgcggctgtcggccgatacatttttggcggcaggcttgggcgcagacctttggcatcgtcaatg | Endogenous plasmid  pRPA |

**Supplementary Table IV.** Segregation primer sets

| Primer Name | Sequence | Amplicon length (bp) | Strain |
| --- | --- | --- | --- |
| upcm_F | GTAGATGACCTTGGC  GAAGAACTTGTCG | 1959 | Wild Type |
| cmKO_R | CGATCCAGACGTC  ATGGCCGATCTC |  |  |
| cmKO_F | TCAACGACTCGCA  GATCACCTACACC | 1999 | Wild Type |
| downcm_R | CCACAACAGATCGCTTTCGCTGC |  |  |
| upcm_F | GTAGATGACCTTGGCGAAGAACTTGTCG | 3743 | Wild Type |
| downcm_R | CCACAACAGATCGCTTTCGCTGC | 3171 | Δcat |
| 25310 KO_F | CCACAAATATCG  GGACTGGTTGG | 2738 | Wild Type |
| 25310 KO_R | CCTATGAAATGAAAG  ACTGCCTTATTACG | 4539 | Δ25310 |
| 24785 KO_F | GTCGATGCGCGT  GACCAGCAGCAC | 2858 | Wild Type |
| 24785 KO_R | CGTTGCCCTTGC  TCACTCTGCACC | 4480 | Δ 24785 |
| 24780 KO_F | GCTTCAGTGACAAAA  GACCAAGCCGCATG | 3014 | Wild Type |
| 24780 KO_R | GGACAAAACCACCCTC  TAACCACCTTGATTC | 4539 | Δ 24780 |
| 25310_F | GCTGCTGGAC  TACGCCATGG | 353 | Wild Type |
| 25310_R | CCTTTGATCCAA  GGCGACACAG |  |  |
| 24785_F | CGTCGTTCCG  AAATAGGTGG | 492 | Wild Type |
| 24785_R | CCATTCCTCAC  CAAAGACAGG |  |  |
| 24780_F | GCAGAACATCT  TTCGAGCCTTC | 514 | Wild Type |
| 24780_R | CGTCTCGCTC  TGCTTTGTTG |  |  |

**Supplementary Table V.** RBS Library

| Colony number | Sequence | Translation Initiation Rate |
| --- | --- | --- |
| 205 | cttggc | 54 |
| 187 | gcatcg | 72 |
| 202 | tgtccg | 81 |
| 392 | gcaagc | 116 |
| 19 | atttga | 167 |
| 232 | acatac | 174 |
| 277 | agatat | 188 |
| 338 | gtggct | 210 |
| 282 | actggt | 244 |
| 224 | caggc | 270 |
| 21 | tggact | 365 |
| 227 | aggcct | 396 |
| 134 | gggctt | 479 |
| 383 | ggtcga | 517 |
| 46 | gtgtag | 622 |
| 444 | tggagc | 665 |
| 437 | tggagc | 665 |
| 239 | gggtca | 680 |
| 45 | aatgag | 1038 |
| 399 | ggtttc | 1081 |
| 197 | gaggct | 1406 |
| 442 | gagggc | 1603 |
| 279 | atggag | 1613 |
| 382 | gtggag | 1613 |
| 108 | gtggag | 1613 |
| 126 | attagg | 2057 |
| 225 | gggtt | 2127 |
| 32 | aggaga | 3029 |
| 159 | aagagg | 3374 |
| 204 | ggggtt | 3700 |
| 226 | acgagg | 4004 |

**Supplementary Table VI.** RT-qPCR primers

| **Primer Name** | **Sequence** | **Amplicon Length (bp)** | **Primer Concentration (nM)** | **cDNA or gDNA Concentration (ng/μL)** | **cDNA or gDNA** | **Efficiency** | **R^2** |
| --- | --- | --- | --- | --- | --- | --- | --- |
| 16SrRNA_F | GCTTAACACAT  GCAAGTCGAAC | 230 | 40 | 0.05 | cDNA | 98.61 | 0.994 |
| 16SrRNA_R2 | TCATCCTCTCA  GACCAGCTAC |  |  |  |  |  |  |
| qmrfp_F | CGTTTCAAAGTTC  GTATGGAAGGTTC | 188 | 50 | 0.25 | cDNA | 104.89 | 0.996 |
| qmrfp_R | GTGTTTAACGTAAG  CTTTGGAACCGTAC |  |  |  |  |  |  |
| qKan_F | CAGTCATAGCCG  AATAGCCTCTC | 161 | 40 | 0.25 | gDNA | 97.64 | 0.979 |
| qKan_R | CAAAGTAAACTGG  ATGGCTTTCTTGC |  |  |  |  |  |  |
| qGent_F | CAAAGTTAGGTG  GCTCAAGTATGG | 182 | 60 | 0.25 | gDNA | 99.04 | 0.968 |
| qGent_R | CGATGAATGTCTT  ACTACGGAGCAAG |  |  |  |  |  |  |
| dxs_F | CTTCTCGACA  CCATTCGCAC | 169 | 40 | 0.25 | gDNA | 95.94 | 0.999 |
| dxs_R | ACATAGTGCAG  TGCGGTAGTC |  |  |  |  |  |  |
| qgfpuv_F | GAGTAAAGGAGAAG  AACTTTTCACTGGAG | 197 | 50 | 40 | cDNA | Just Visualized | |
| qgfpuv_R | CCATAAGAGAAAGT  AGTGACAAGTGTTG |  |  |  |  |  |  |
| qCAT_F | GCACCATCCTG  AACGAAGTC | 187 | 50 | 40 | cDNA | Just Visualized | |
| qCAT_R | GCCTTCGAAA  TAGGTGCTGG |  |  |  |  |  |  |
| qRepA_F | GCACGAAGAA  CATGAGCTGC | 178 | 50 | 0.25 | gDNA | 94.71 | 0.978 |
| qRepA_R | CGACCTTGAGC  TTGAGGGAAAG |  |  |  |  |  |  |

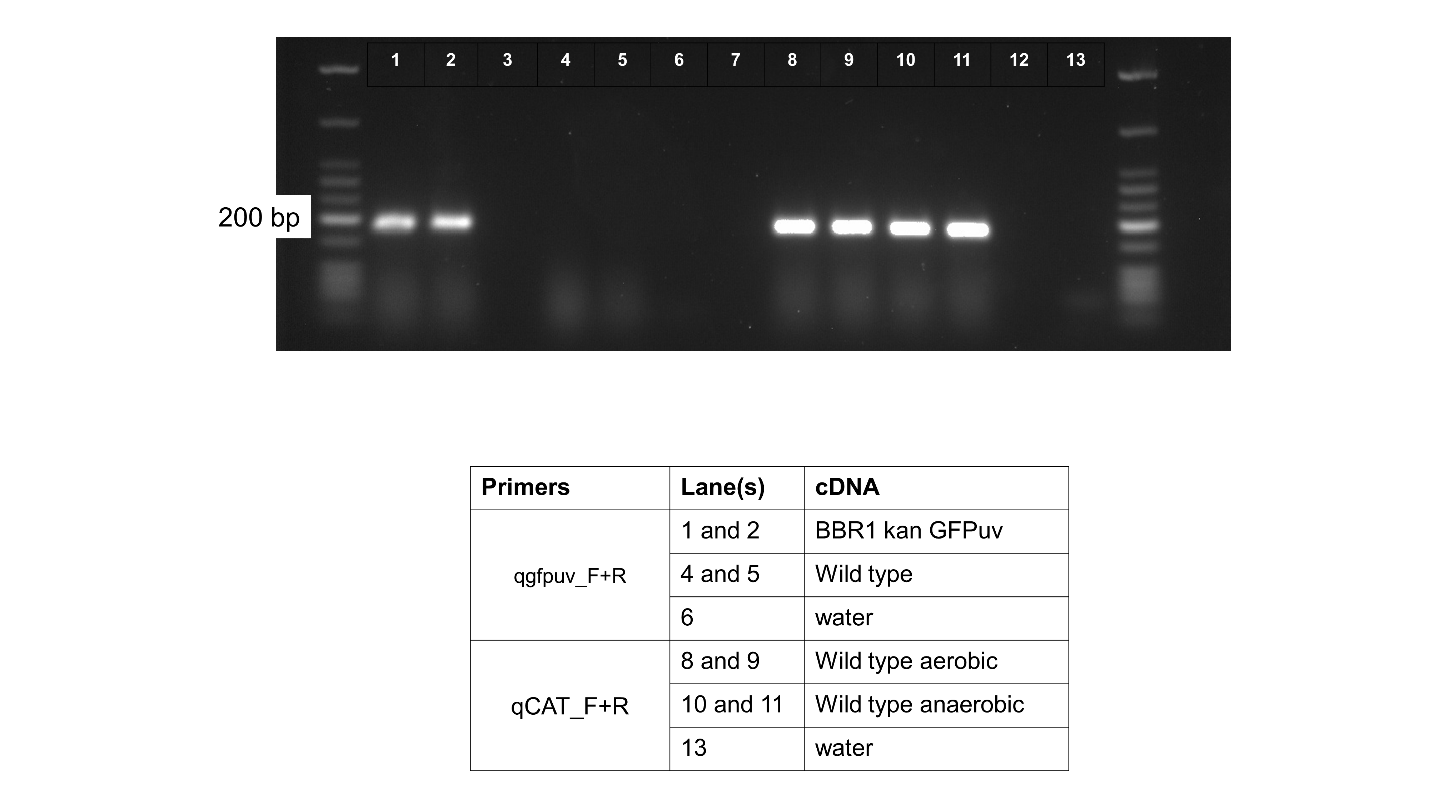

**Supplementary Figure 1.** 2% agarose gel of RT-PCR products showing expression of *gfpuv* in the BBR1 kan GFPuv strain but not the wild type, and expression of *cat* in the wild type strain grown aerobically and anaerobically.

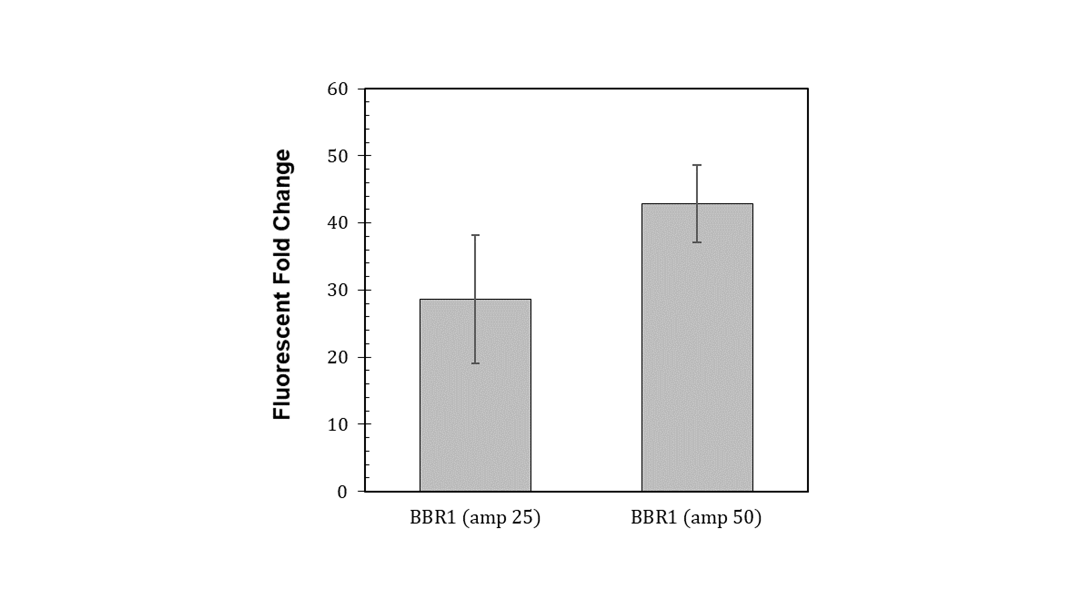

**Supplementary Figure 2.** Fluorescent fold change average of *R. palustris* BBR1 amp mRFP strain grown in triplicate with 25µg/mL or 50 µg/mL ampicillin added to the media (Materials and Methods). Error bars indicate one standard deviation.

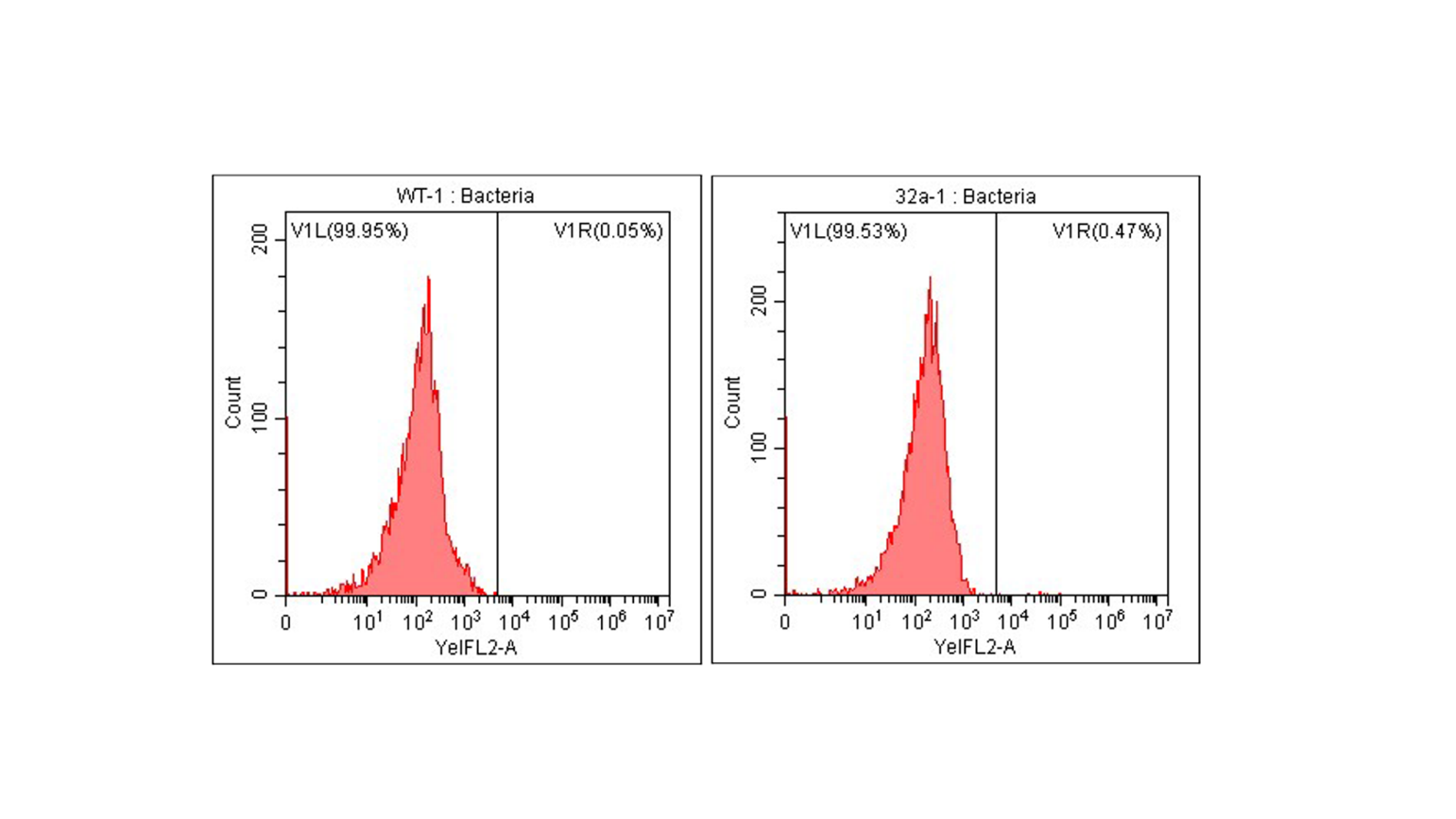
**Supplementary Figure 3.** Representative flow cytometry results of mRFP fluorescence. **A)** Wild type *R. palustris* **B)** *R. palustris* BBR1 amp mRFP strain grown with 100 µg/mL ampicillin added to the media (Materials and Methods).

**A)**

**B)**

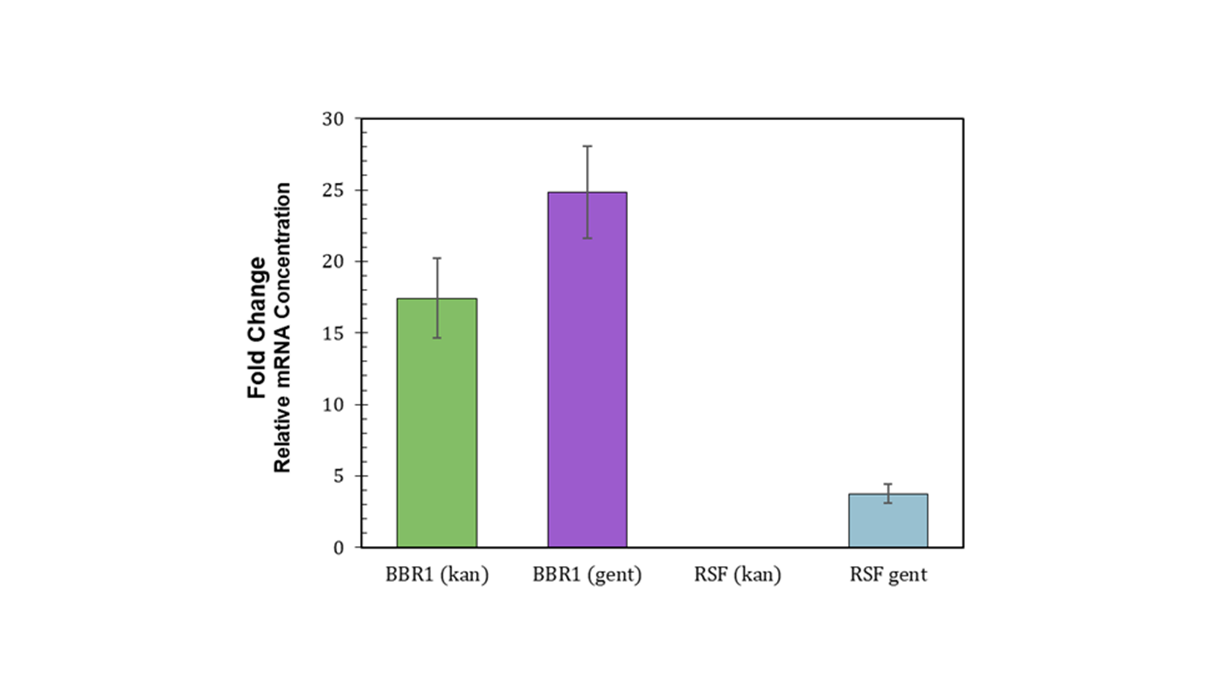

**Supplementary Figure 4.** Relative mRNA concentration (*mrfp* relative to *16rRNA*) for the *R. palustris* BBR1 kan, BBR1 gent, RSF1010 kan, and RSF1010 gent strains respectively (Materials and Methods). Two biological and two technical replicates were averaged for each strain. Error bars indicate one standard deviation.
